# First computational design of Covid-19 coronavirus vaccine using lambda superstrings

**DOI:** 10.1101/2020.11.30.403824

**Authors:** Luis Martínez, Iker Malaina, David Salcines, Héctor Terán, Santos Alegre, I.M. De la Fuente, Elena Gonzalez Lopez, Gonzalo Ocejo Vinyals, Carmen Álvarez

## Abstract

In this work we have developed, by employing lambda superstrings, a map of candidate vaccines against SARS-CoV-2 with lengths between 9 and 200, based on estimations of the immunogenicity of the epitopes and the binding affinity of epitopes to MHC class I molecules using tools from the IEDB Analysis Resource, as well as the overall predictions obtained using the VaxiJen tool. We have synthesized one of the peptides, specifically the one of length 22, and we have carried out an immunogenicity assay and a cytokine assay, which has given positive results in both cases.

## 1. Introduction

The Covid-19 coronavirus epidemic represents the greatest global threat to human health at the current juncture, with more than 60 million persons infected and more than 1.4 million persons died worldwide in the 11 months since the disease was detected. In order to stop this epidemic, different types of vaccines are being developed in an accelerated manner, mostly, if not all, based on well-known classical methods. Despite the fact that several effective vaccines are available, their effectiveness has been demonstrated only in clinical trials, and it remains to be checked if the obtained percentages of protection will be maintained in the whole population and, above all, for how long will the obtained immunity last. This makes imperative to continue searching for new candidate vaccines against SARS-CoV-2.

We present here a candidate for a vaccine for the SARS-CoV-2 virus designed entirely from computational methods and advanced tools of artificial intelligence. This new method of intelligent vaccine design is based on developing, first, one or several candidates that allow to cover the maximum number of possible mutations, doing then the molecular sequencing of the candidates, and performing then the subsequent in vitro and in vivo tests and, finally, the clinical analysis.

Our method is based in the concept of λ-superstring, introduced by our research team in [6]. In that paper, we presented a new criterion for the selection of epitopes in the design of vaccines which is well suited to consider all the mutations of the pathogen so that there is a balance of the covering of the candidate vaccine among all the mutated versions of the target protein.

Specifically, we considered a set of target strings, formed by the epitopes that can be selected for the candidate vaccine, and a set of host strings, constituted by the different variants of the target protein, in which the known mutations are considered. In that context, given the value of a parameter λ, a λ-superstring is a sequence of aminoacids with the property that the string covers at least λ epitopes in each of the host strings.

The concept of λ-superstring was generalized later by us in [7] to the one of weighted λ-superstring by allowing the epitopes to be weighted by their immunogenicities. This generalization entails an important improvement in the applications to vaccine design, as this weighting of the immunogenicities gets closer to the biological and medical practice, and increases the likelihood that the obtained candidate vaccines are effective. In fact, the use of weighted λ-superstrings could be useful to fight the high mutability and escape mutations of viruses like HIV, HCV, or Influenza.

Here, we have used Integer Programming to obtain the best solution for the weighted lambda-superstring problem, applied to the surface protein of SARS-COV-2, obtaining sequences for the Spike protein which offer potential protection against all the virus variants considered for the study (in this case, all the sequences appearing in GISAID ([4]) webside (until March 4, 2020). In consequence, the objective of our proposal is not only to develop an effective vaccine for the current pandemic, but also to give protection against potential mutations of the virus, and stop its expansion and possible comeback.

Specifically, in order to select the most promising epitopes for the vaccine, we have first associated a weight to each potential 9-mer epitope. More precisely, we have estimated the immunogenicity and HLA-binding affinity of class I potential epitopes, which makes our candidates appropriate to be used worldwide. As a result of our computations, we present here a map of optimal candidate vaccines with different lengths, which on one hand optimize both immunogenicity and HLA-binding affinity, and on the other hand, give a balanced protection against all the sequenced variants of Covid-19 surface protein so far. We have selected from this map a peptide of length 22 and we have made in-vivo experimental tests that have given positive results in both immunogenicity assays and cytokine assays.

Our approach gives optimal solutions and therefore represents an efficient alternate insight to vaccine design for SARS-CoV-2 virus.

The main objective of this manuscript is that vaccine specialists become aware of this new method of vaccine design, so that, together, as soon as possible, an efficient vaccine against Covid-19 can be developed.

## 2. Materials and Methods

### 2.1 Extraction of the sequences

The sequences were taken from two sources: GenBank ([3]) and GISAID ([4]).

In Genbank the search “Severe acute respiratory síndrome coronavuirus 2” AND “Homo sapiens” was done in Nucleotide. The results of the search were saved as Coding Sequences in format FASTA Protein in order to get the information about the corresponding aminoacids. After that, the generated file was read and the sequences corresponding to “surface” or to “spike” protein (it is the same protein, but in GenBank it appeared as surface in some cases and as spike in other cases) were selected.

The search in GISAID was restricted by selecting “Human” in the host window and clicking “complete” (>29000bp) in order to obtain only complete genomes in human hosts. The information about the surface protein was extracted from the genomes by using the GeneWise ([5]) tool, by putting in the protein window the sequence of aminoacids corresponding to the surface glycoprotein (YP_009724390.1) product in the reference genome (NC_045512.2).

Duplicated sequences were removed, keeping just a single copy of each one of them, as well as the ones containing some of the ambiguous characters “X”,“B”,“Z”,“J”,“O”,“U”,“.”,“*”. An anomalously short sequence of length 35 was disregarded.

The resulting 22 sequences were taken as host strings, constituting the set *H*.

### 2.2 Weighting of the epitopes

The set *T* of target strings was taken to be the set of 9-tuples of elements of *A* (where *A* is the set of 20 aminoacids) that are contained in at least one target string.

The way in which the weight *w*(*s*) associated to a host string (epitope) *s* was calculated is as follows:

1. The estimation *i*(*s*) of the immunogenicity of *s* was calculated with the “T cell class I PMHC immunogenicity predictor” of IEDB.
2. The set *AI*(*s*) of alleles of the HLA-I allele reference set with the “Peptide binding to MHC class I molecules” tool of IEDB which pass the threshold was computed, and the number *bI*(*s*) = Σ_*i*∈*AI*(*s*)_*f*(*a*) was calculated, where *f*(*a*) is the estimated global frequency of the allele *a* in “The Allele Frequency Net Database”.

Next, the normalized families were computed in the following way:

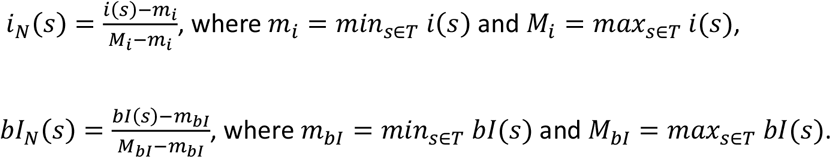

Finally, the weight of the epitope *s* was taken as

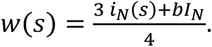

### 2.3 Optimization with CPLEX

CPLEX Optimizer ([2]) was used to solve the integer programming algorithm described in [7] in order to maximize the value of λ for a fixed maximum length with the set of host strings described in Subsection 2.1 and the set of target strings and weights described in Subsection 2.2.

### 2.4 Ranking the candidates with Vaxijen

The bioinformatic tool Vaxijen ([10]) was used with each one of the candidates obtained with the optimization done with CPLEX. “Virus” was selected as target organism. The overall prediction for the protective antigen was calculated for each sequence and the ones over the threshold of 0.4 established for the model were selected.

## 3. Results

Given two sets *H* and *T* of strings, called host strings and target strings, respectively, and given a mapping : *w*: *H* → ℝ, we say that a string *s* is a weighted λ–superstring if, for every *h* ∈ *H*, the inequality Σ_*t*∈*T*,*t*≤*s*_*w*(*t*) ≥ *λ* holds.

We have taken as host strings the 22 distinct sequences corresponding to the Surface protein of SARS-COV-2 appearing in Genbank ([3]) and in the GISAID database ([4]).

We have taken as target strings, that is, as potential epitopes, the 9-mers contained in some of the 22 host strings.

We have weighted the epitopes having into account the estimation of their immunogenicities and the estimation of the binding affinity to HLA-I.

We have calculated a weighted λ–superstring with maximum λ for a length between 9 and 200. For each one of them we have calculated also the VaxiJen overall prediction. These optimal weighted λ–superstring, as well as the corresponding λ values and VaxiJen predictions are shown in Table 1. Each λ–superstring can be divided in a natural way as a union of a small number of peptides located at different parts of the protein. These peptides are enumerated, for each λ–superstring, in the fourth column of the table. When a peptide has some intersection with a domain of the protein, the domain is annotated next to the peptide. For the λ–superstrings of lengths from 176 to 183 the third peptide intersects two domains: NTD and RBD.

**Table 1:**
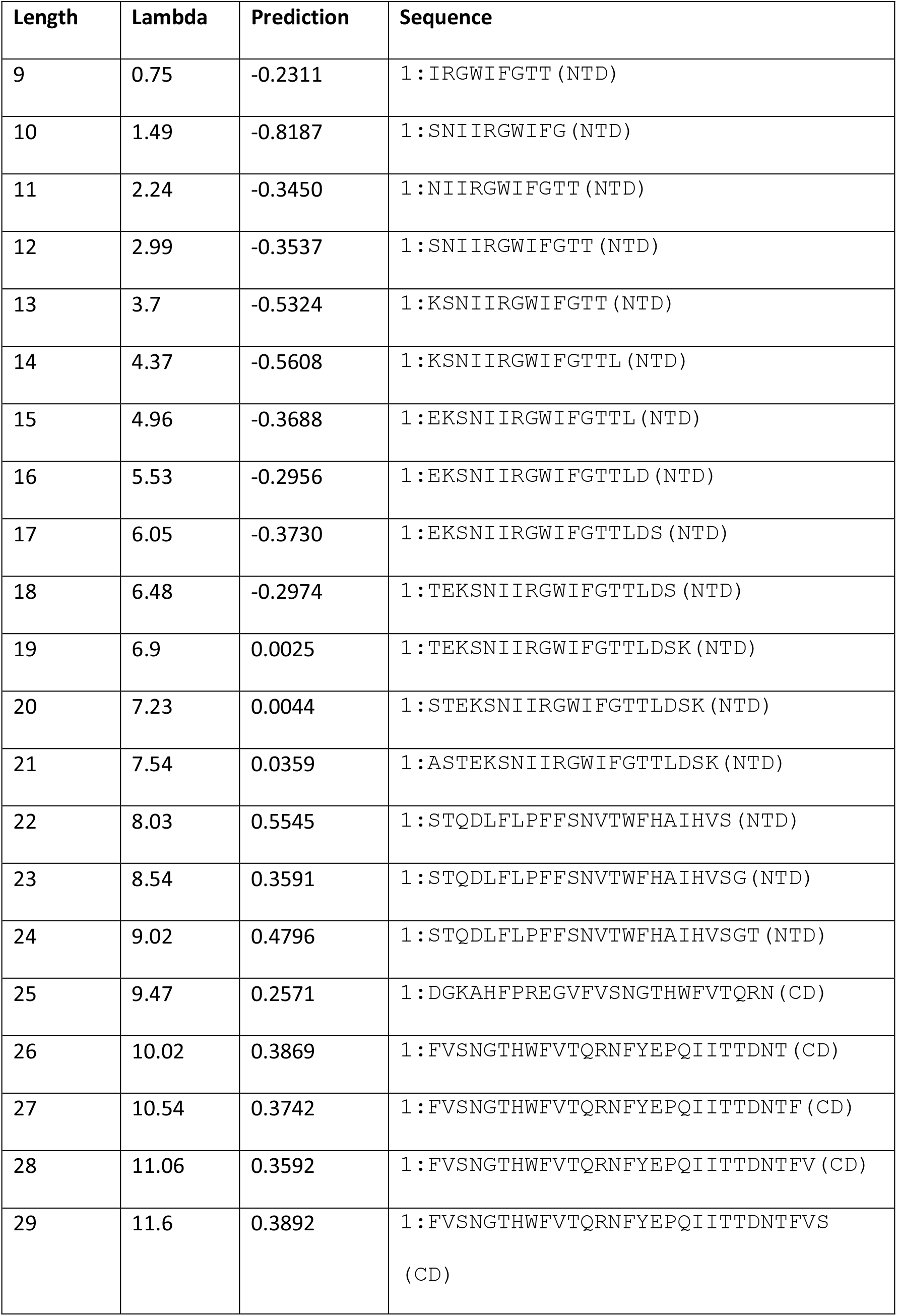
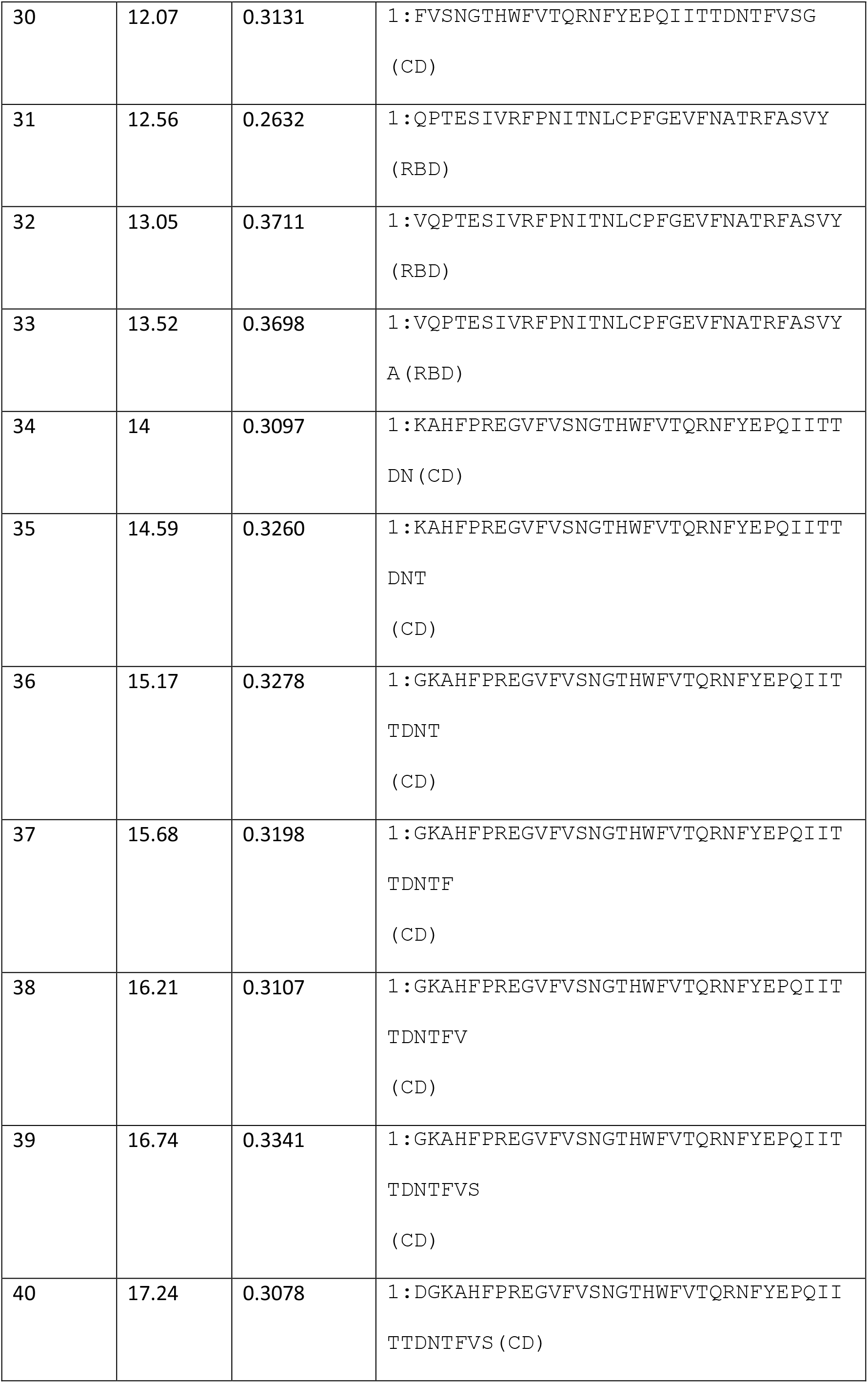
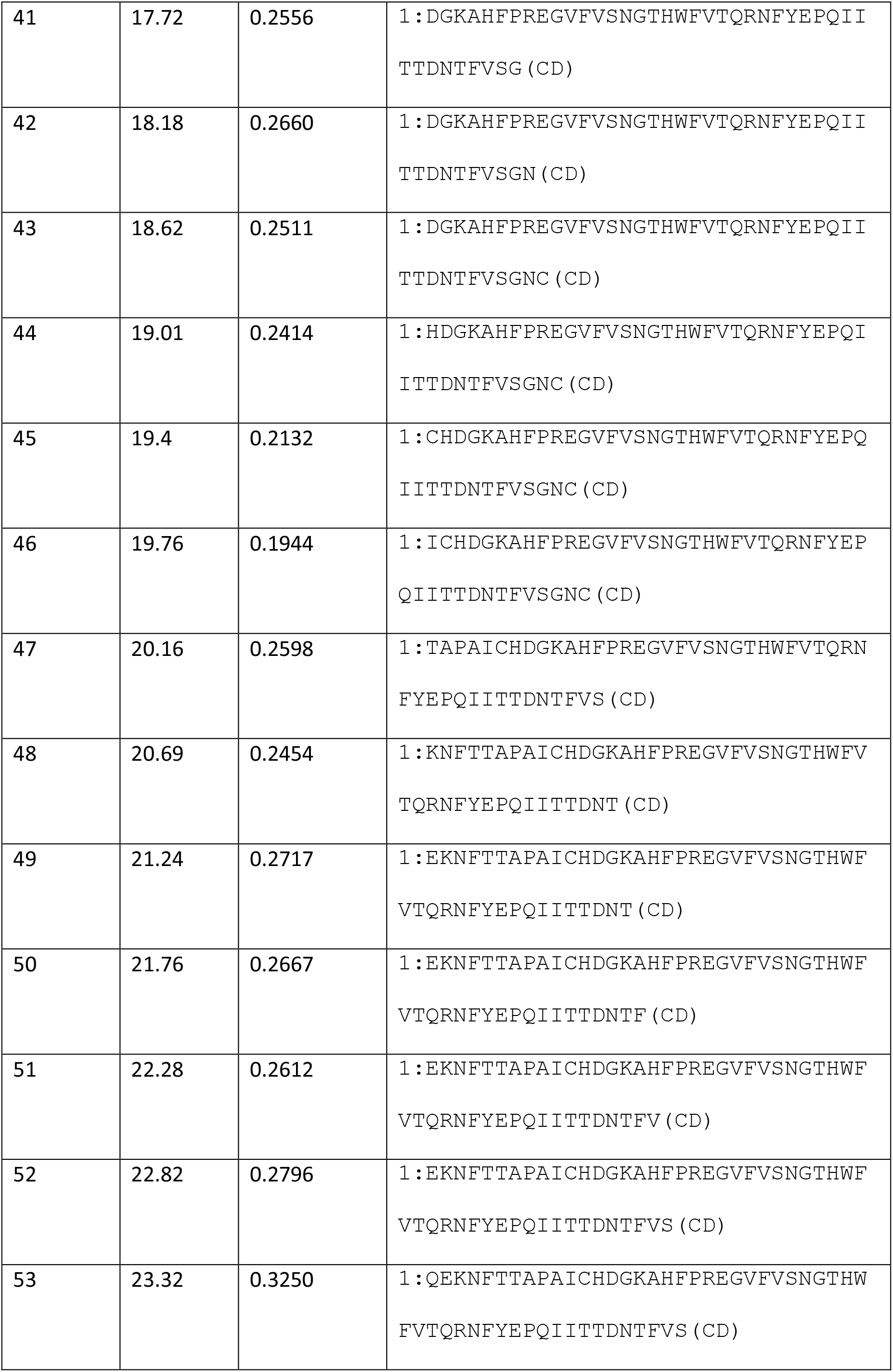
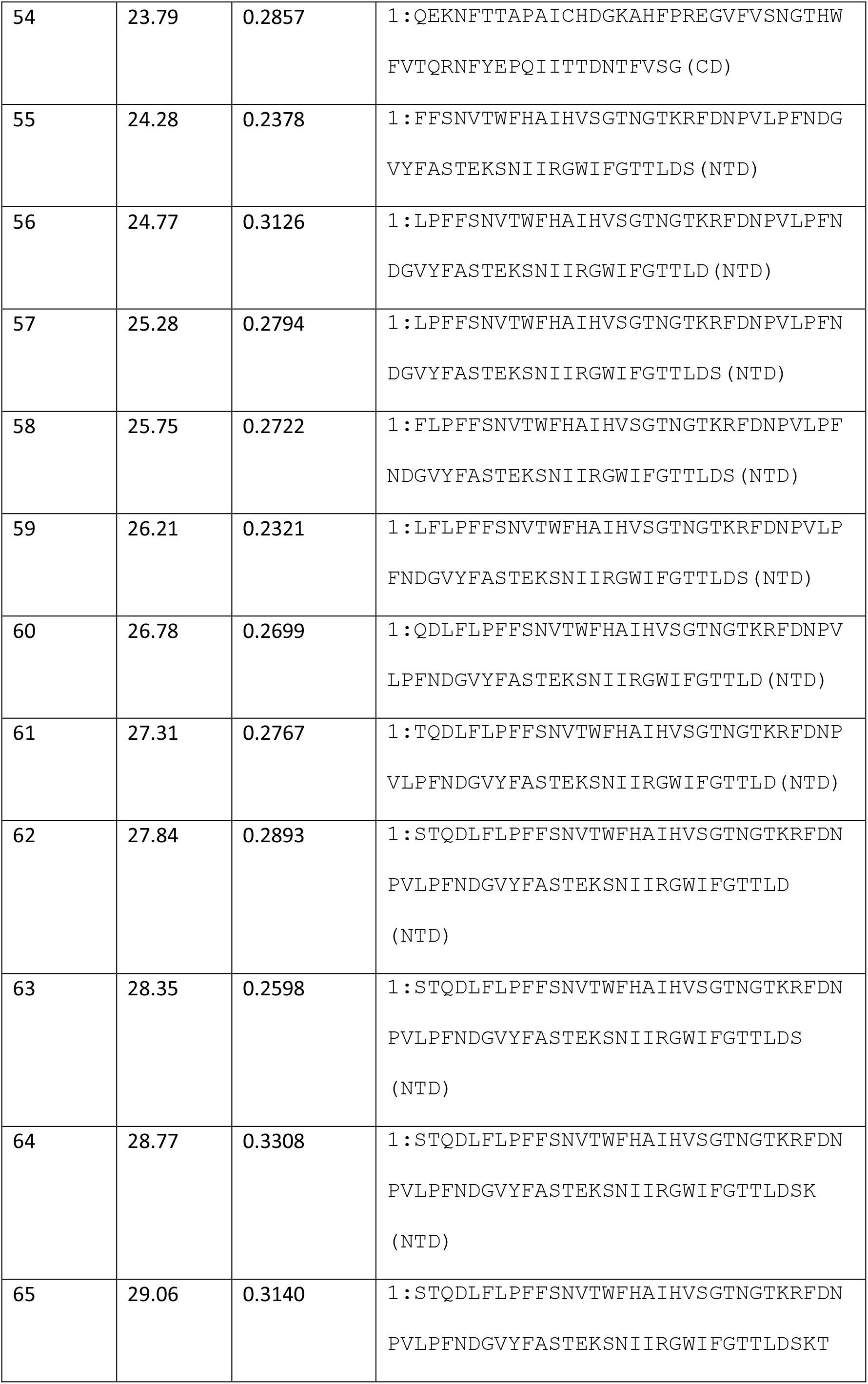
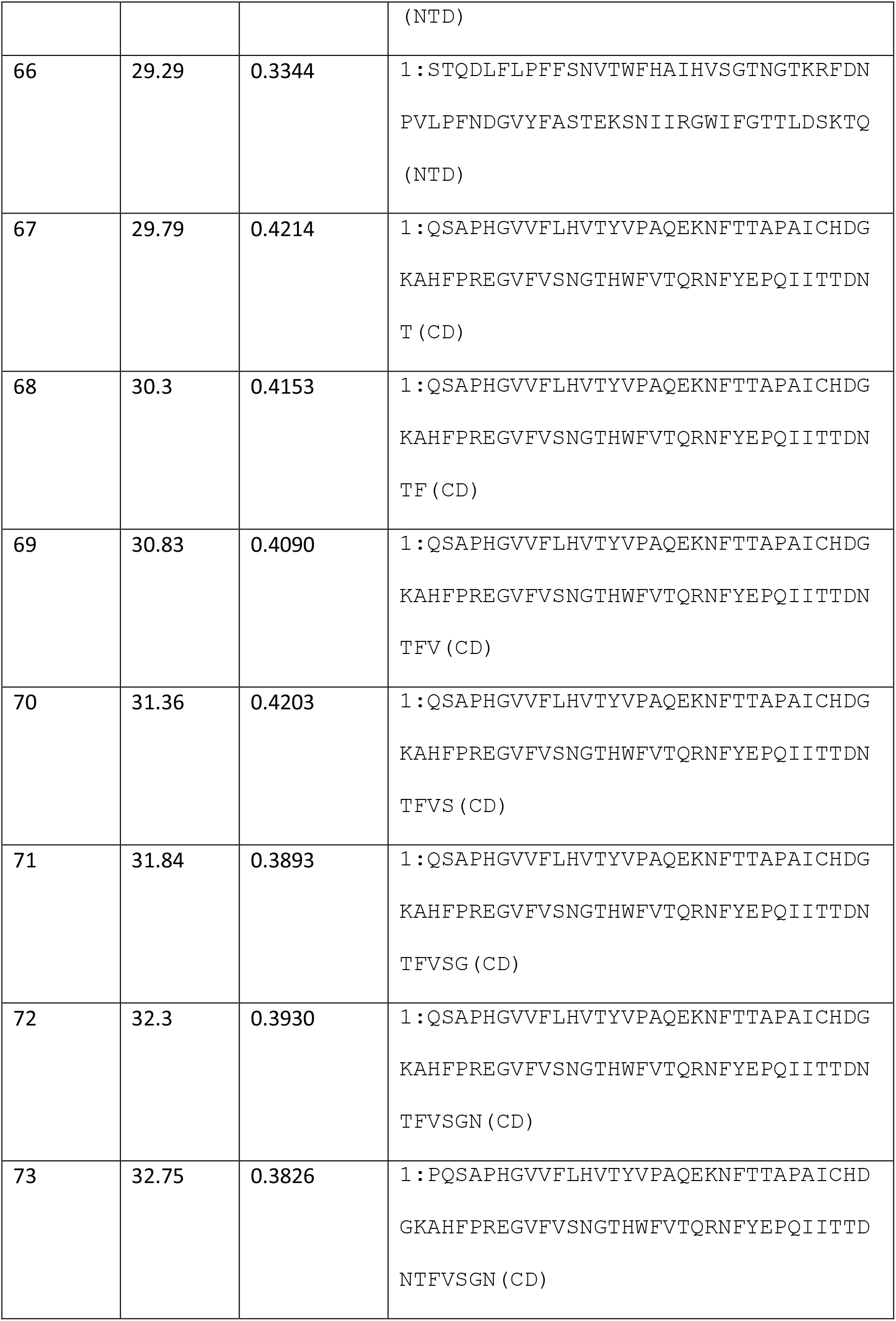
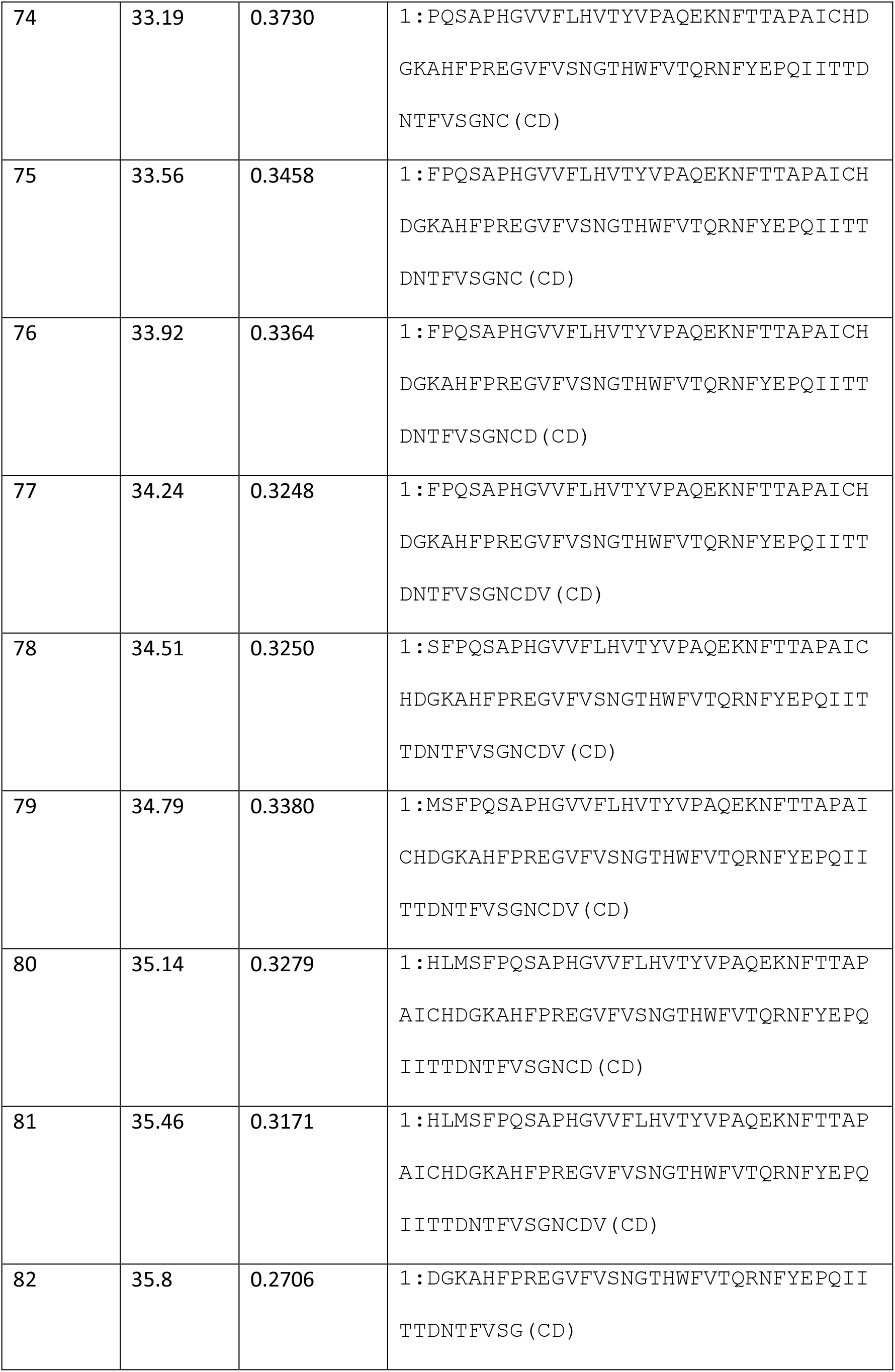
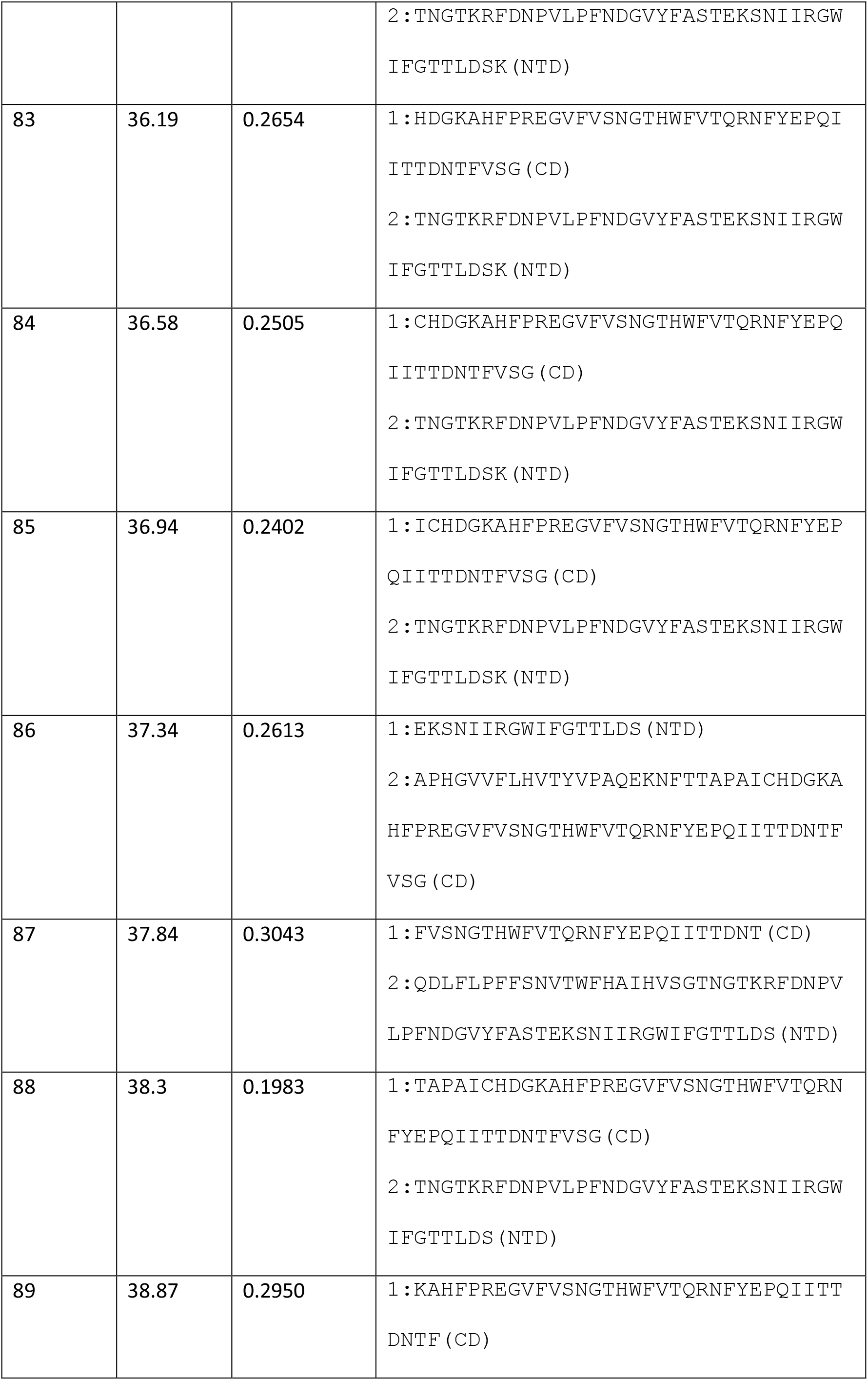
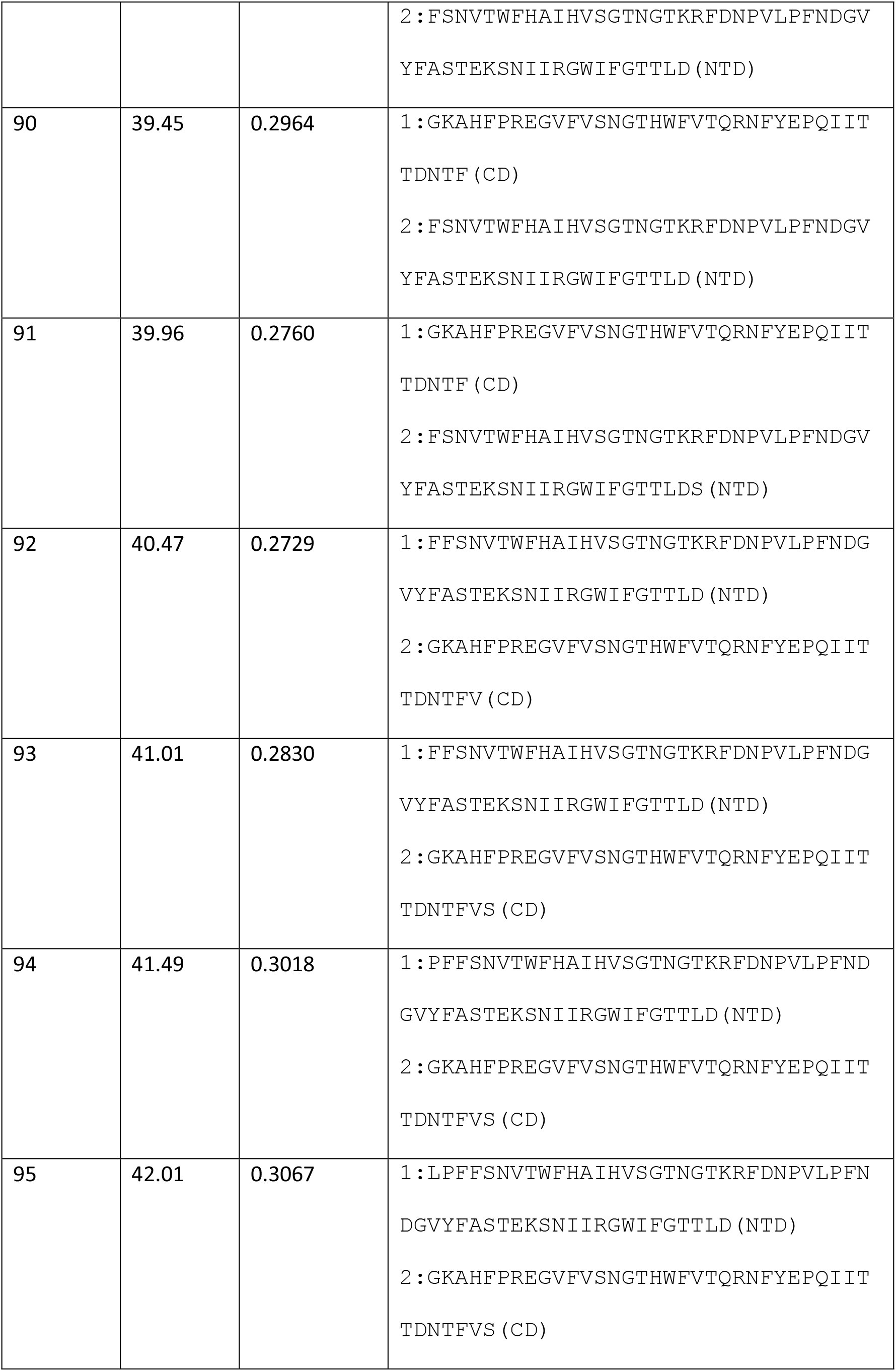
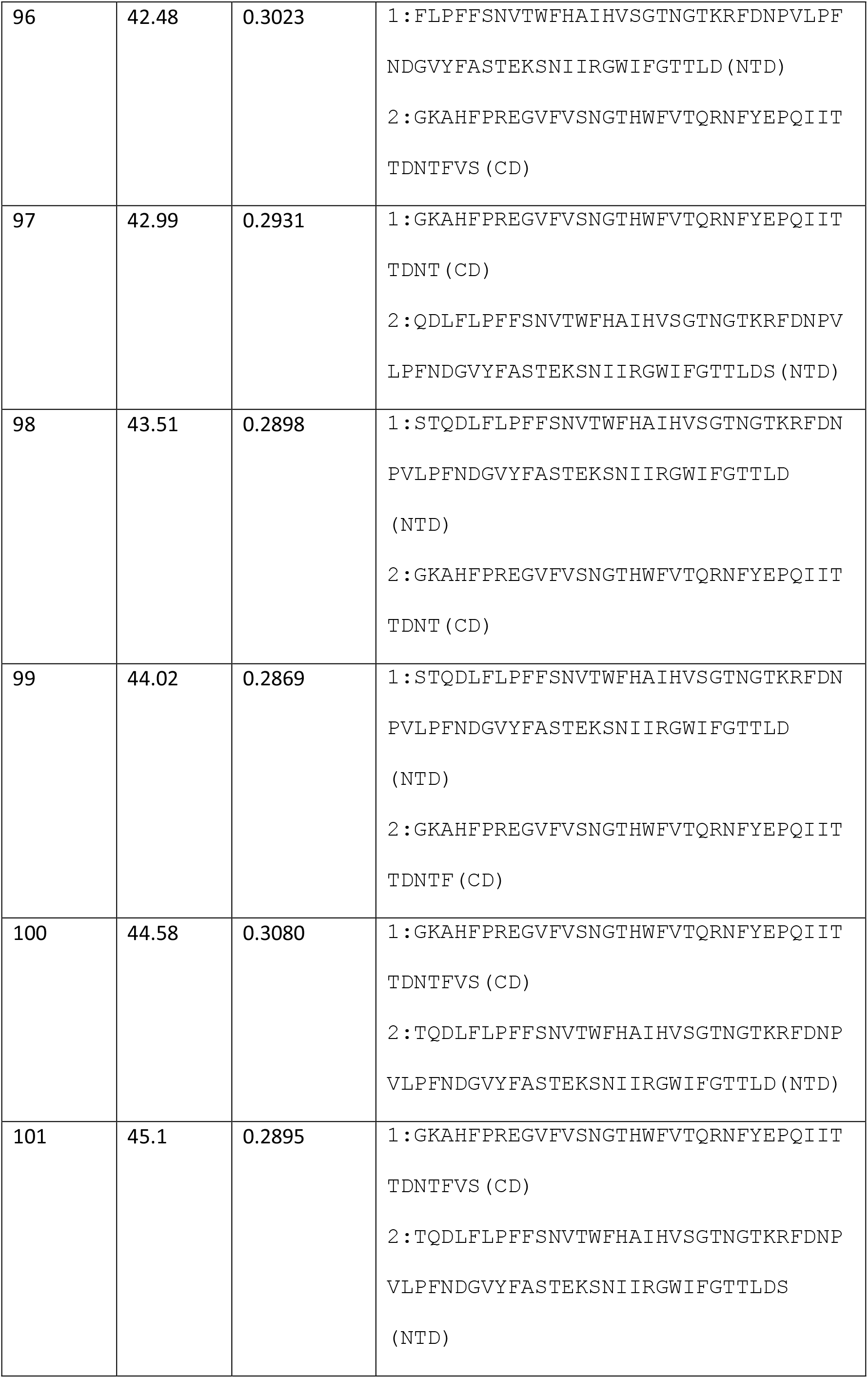
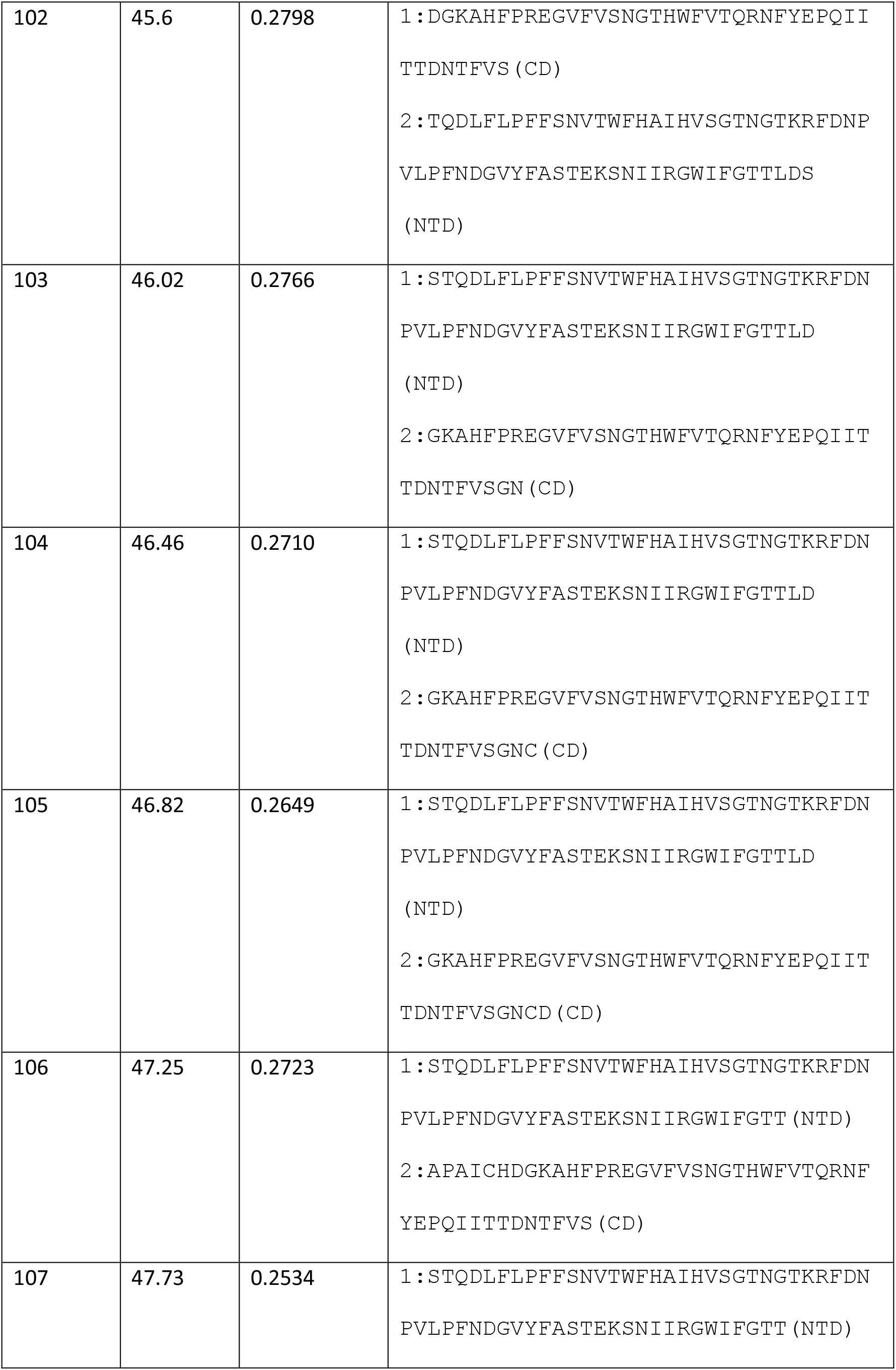
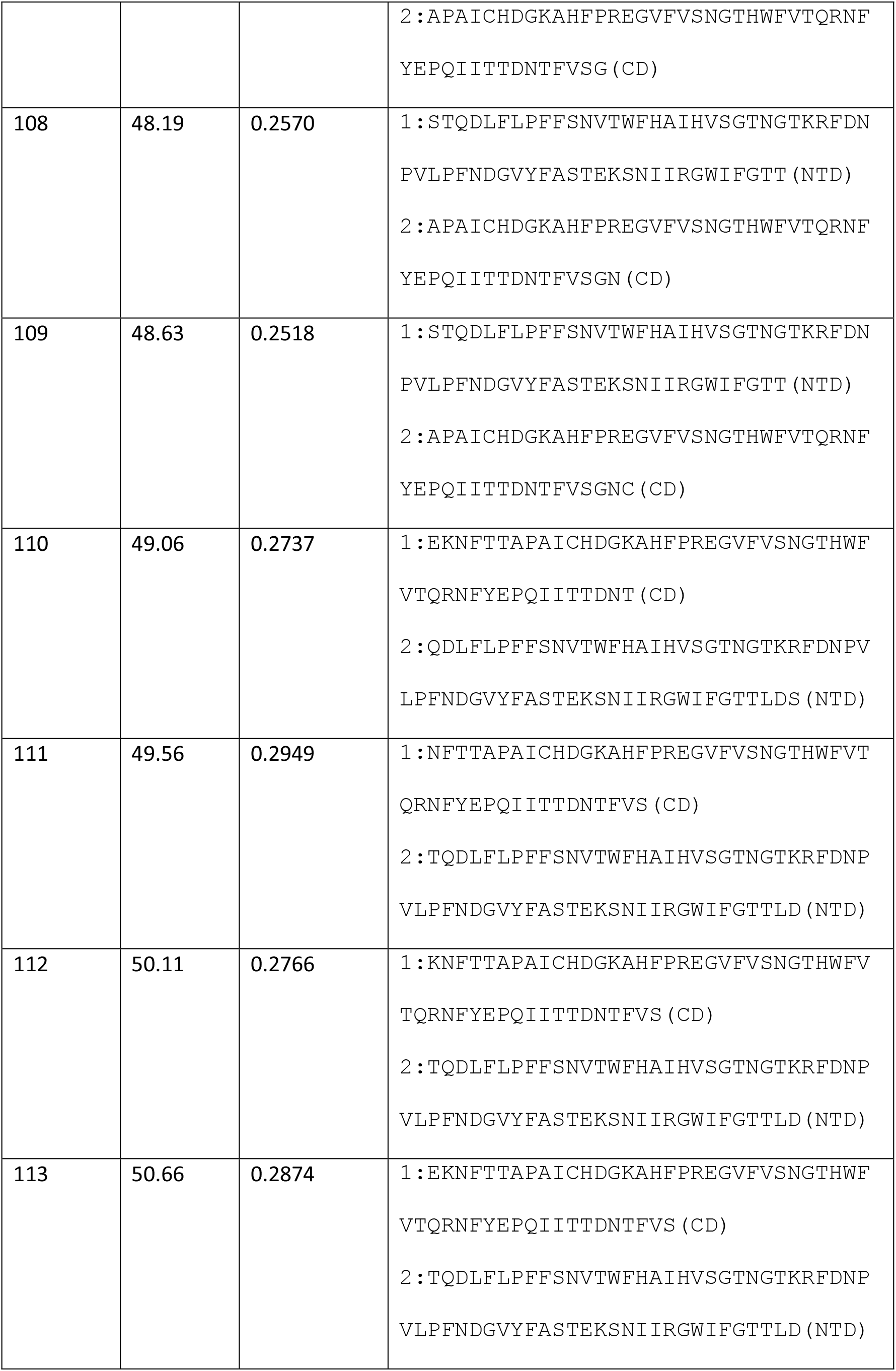
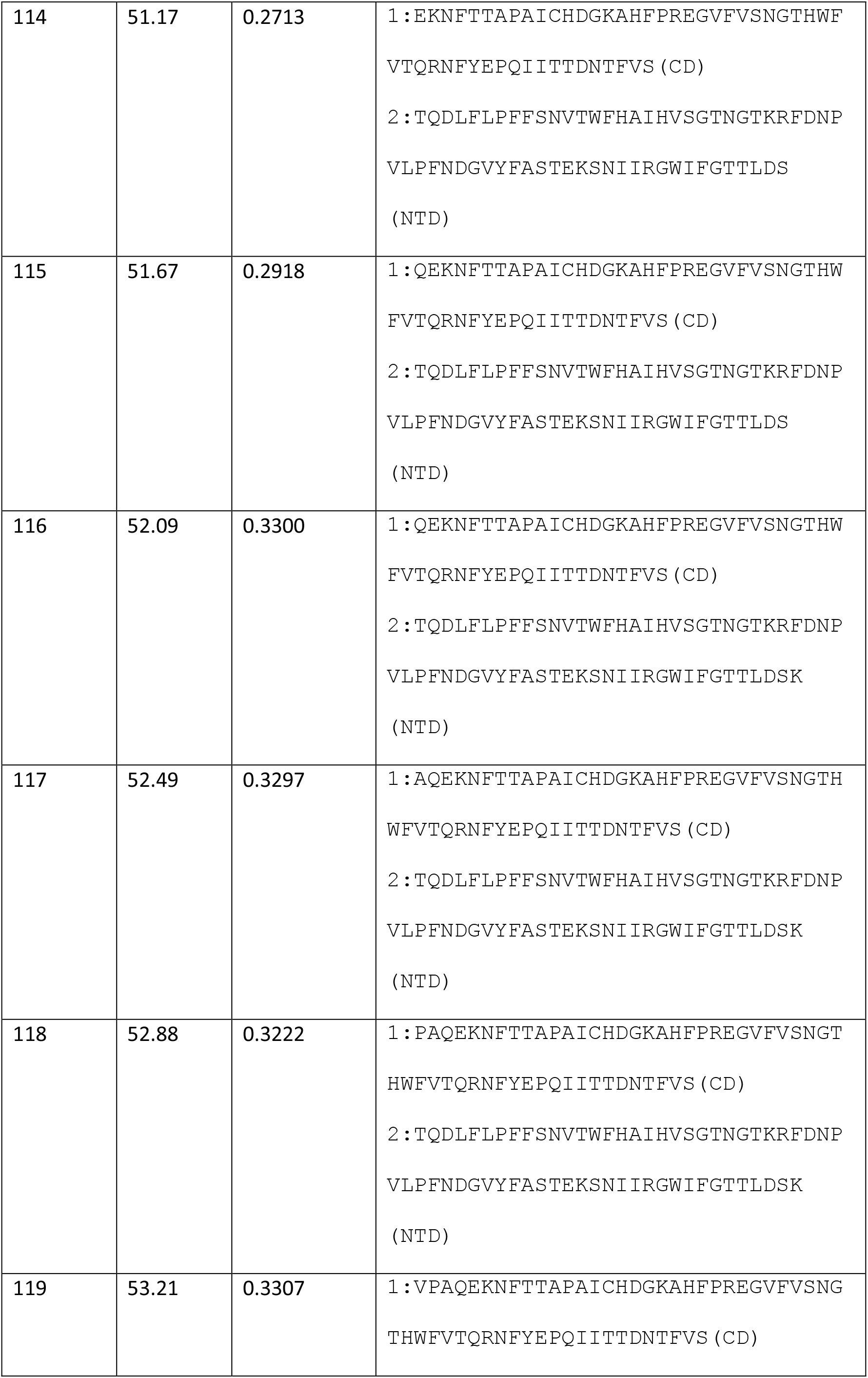
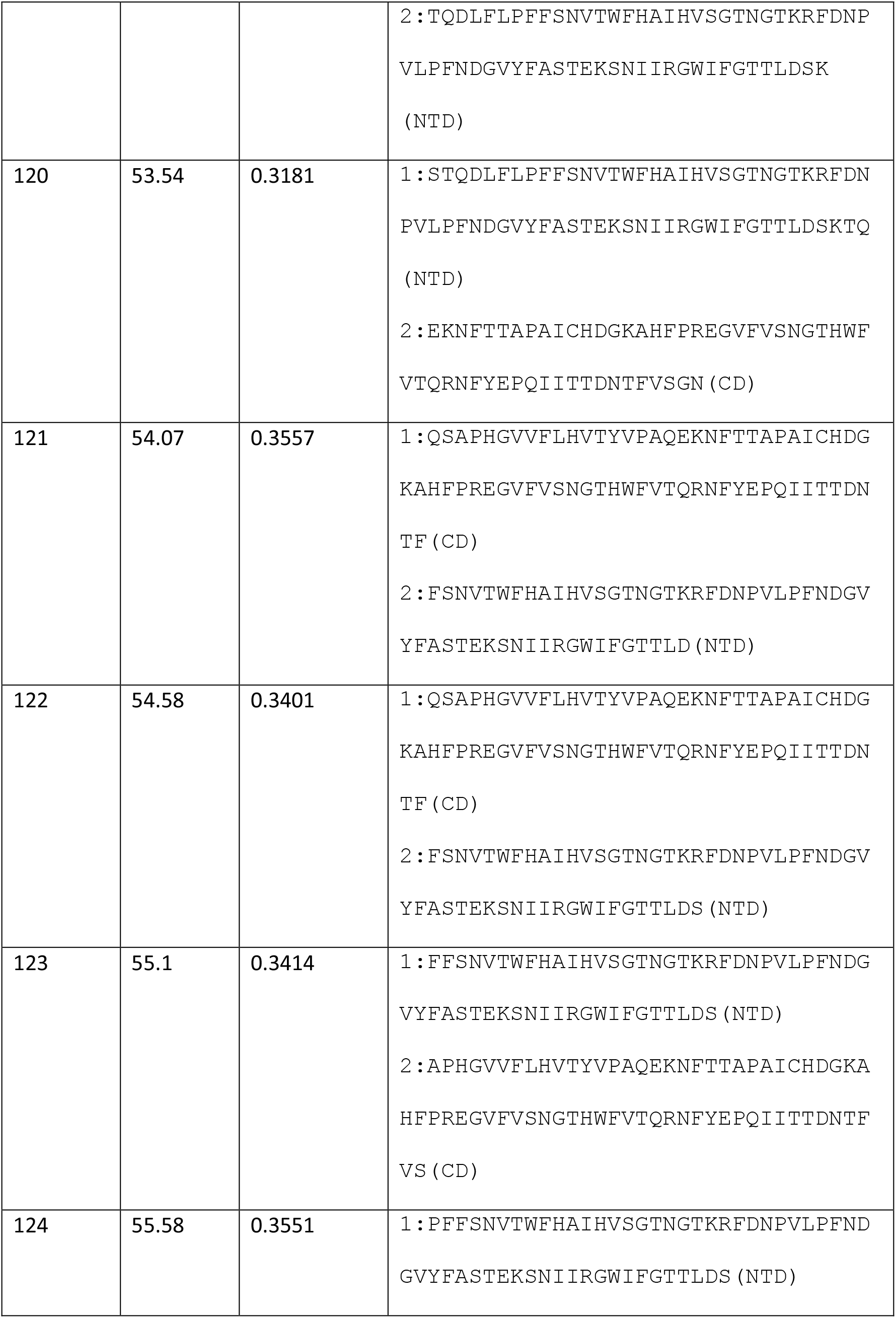
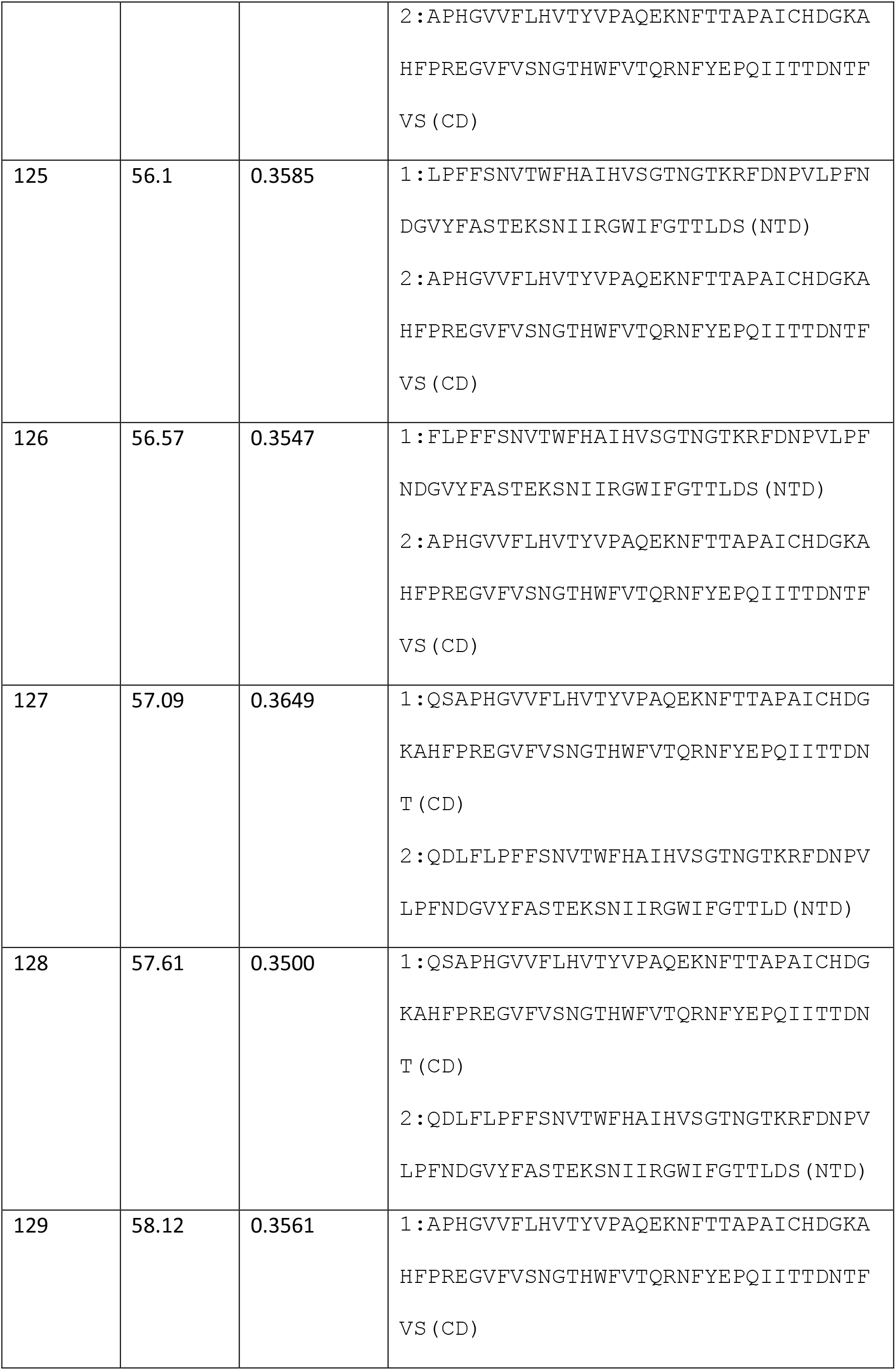
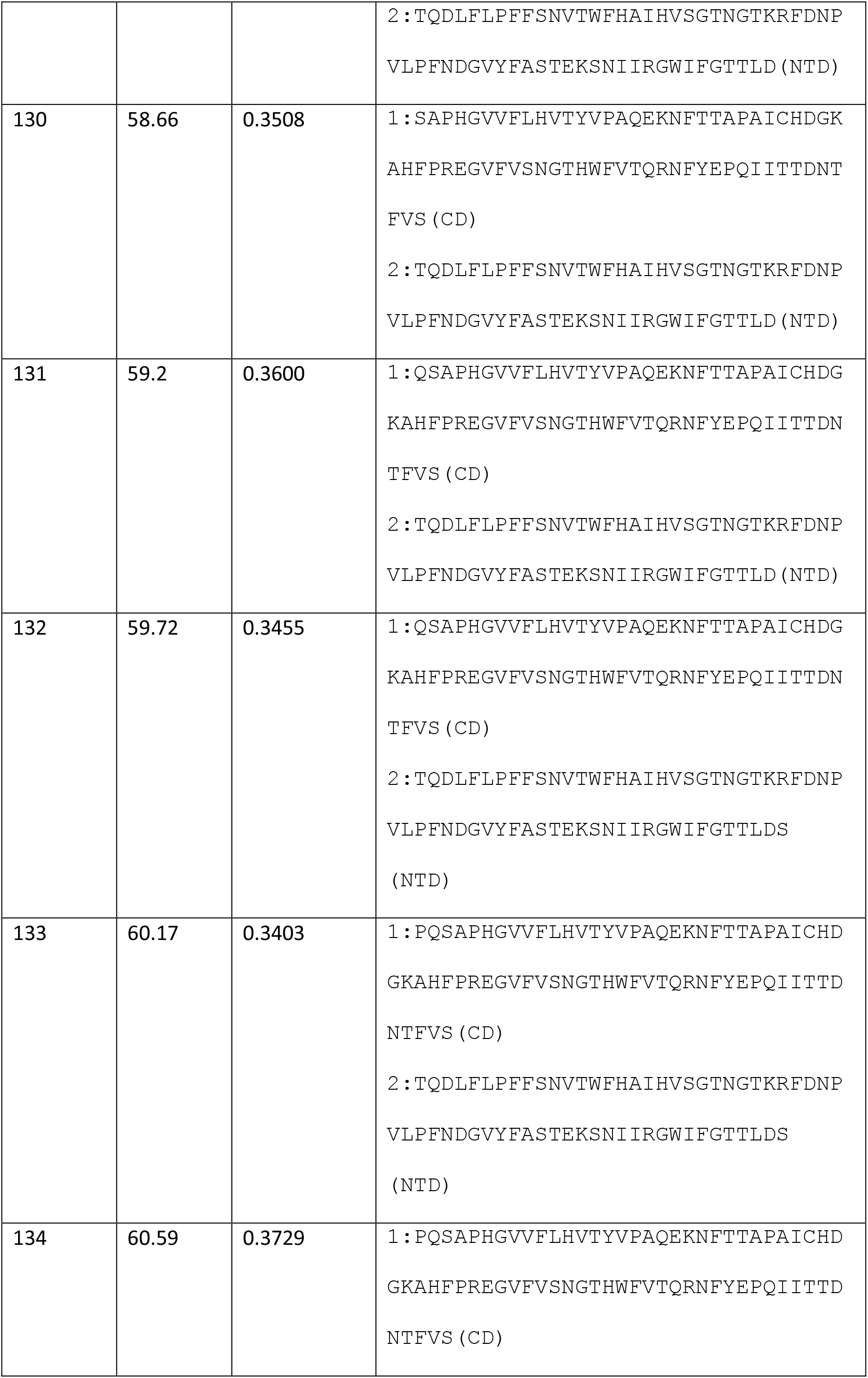
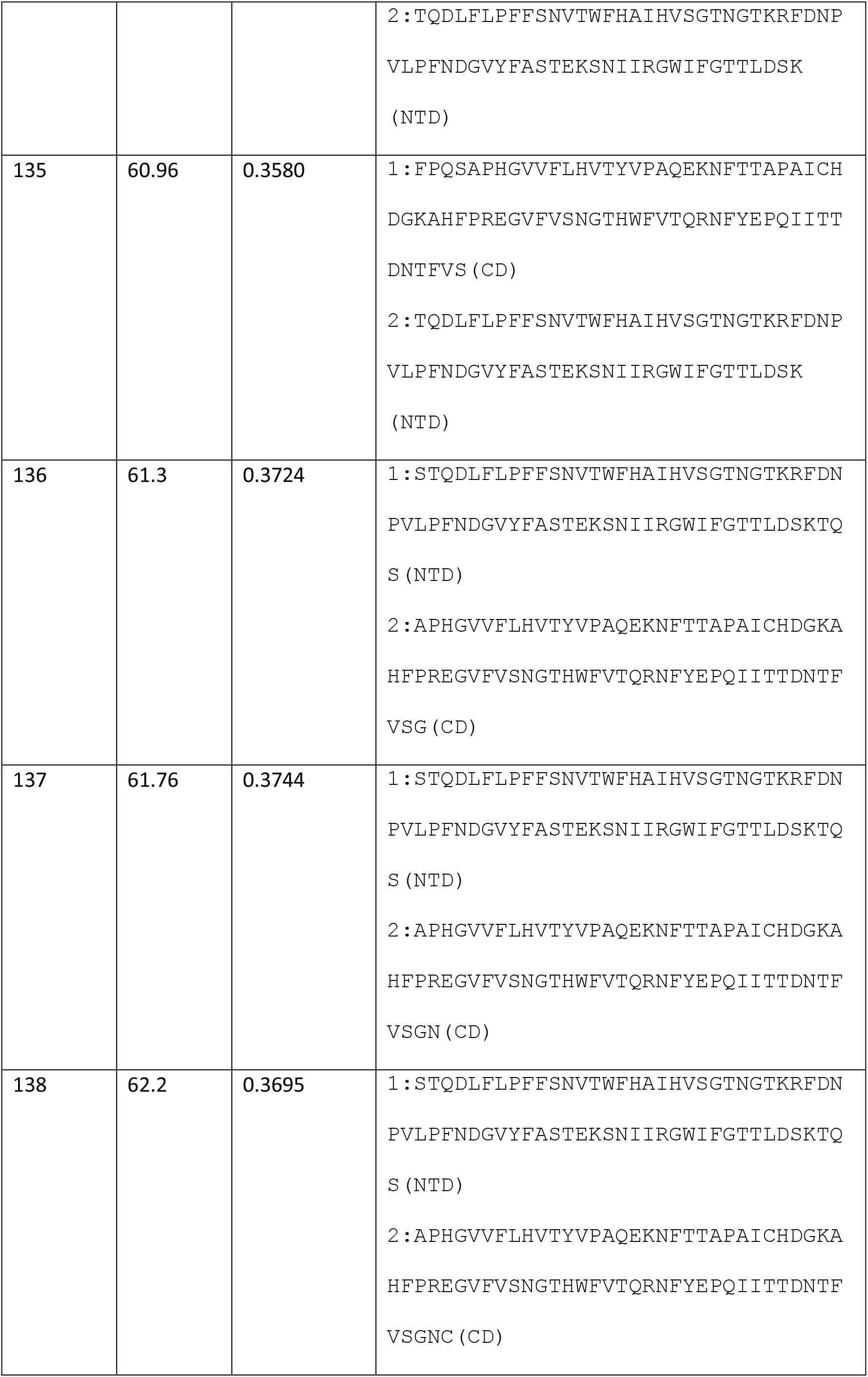
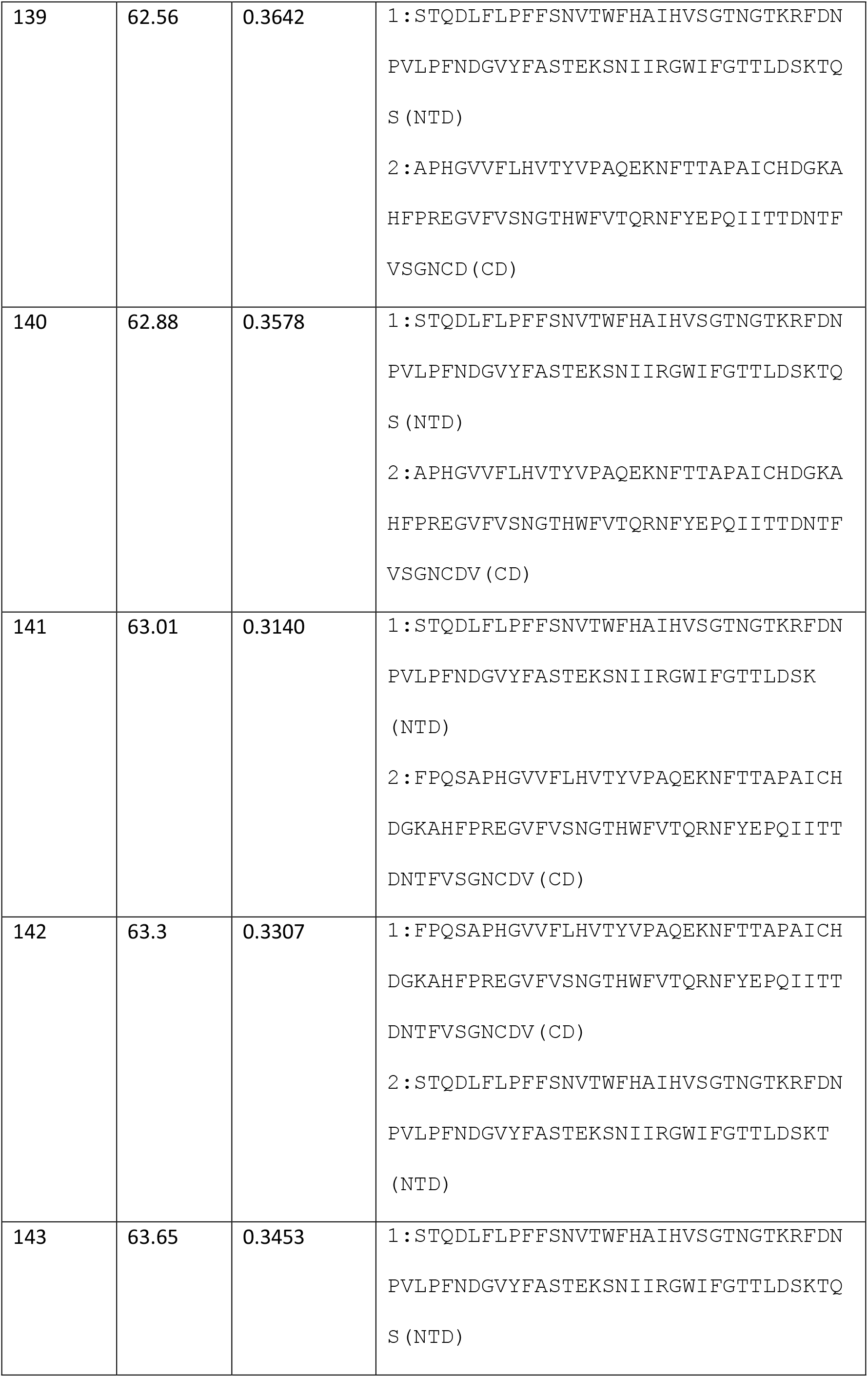
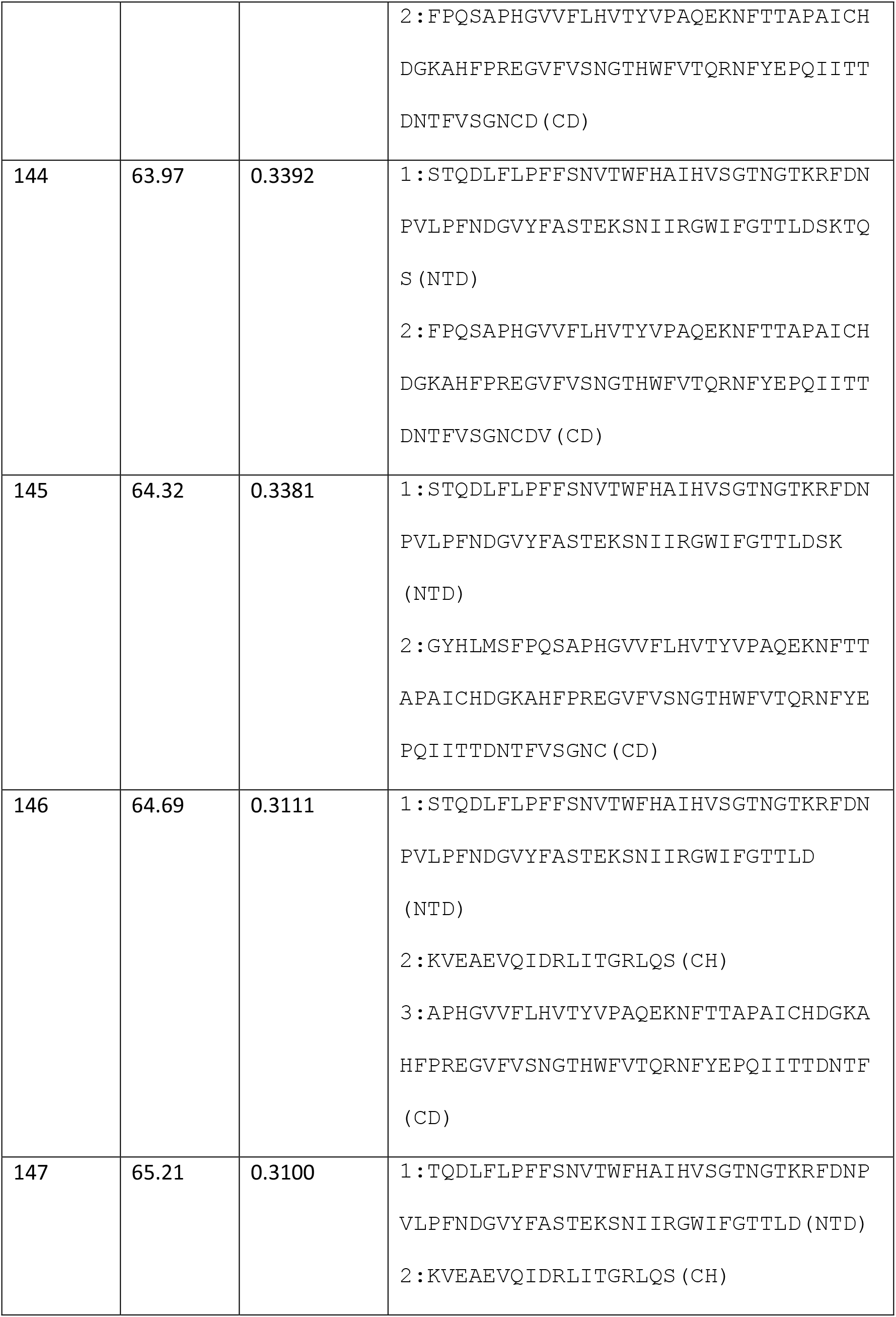
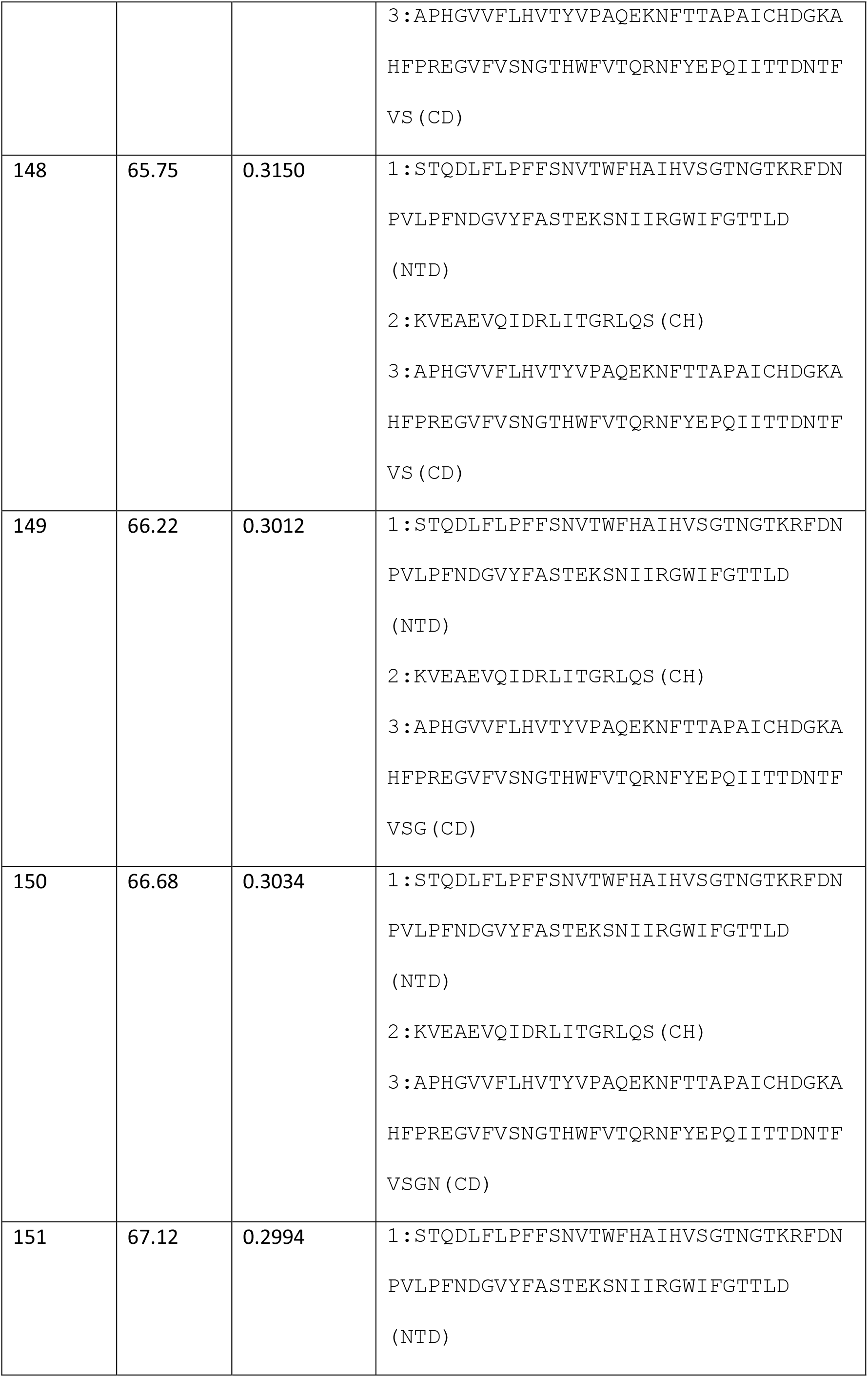
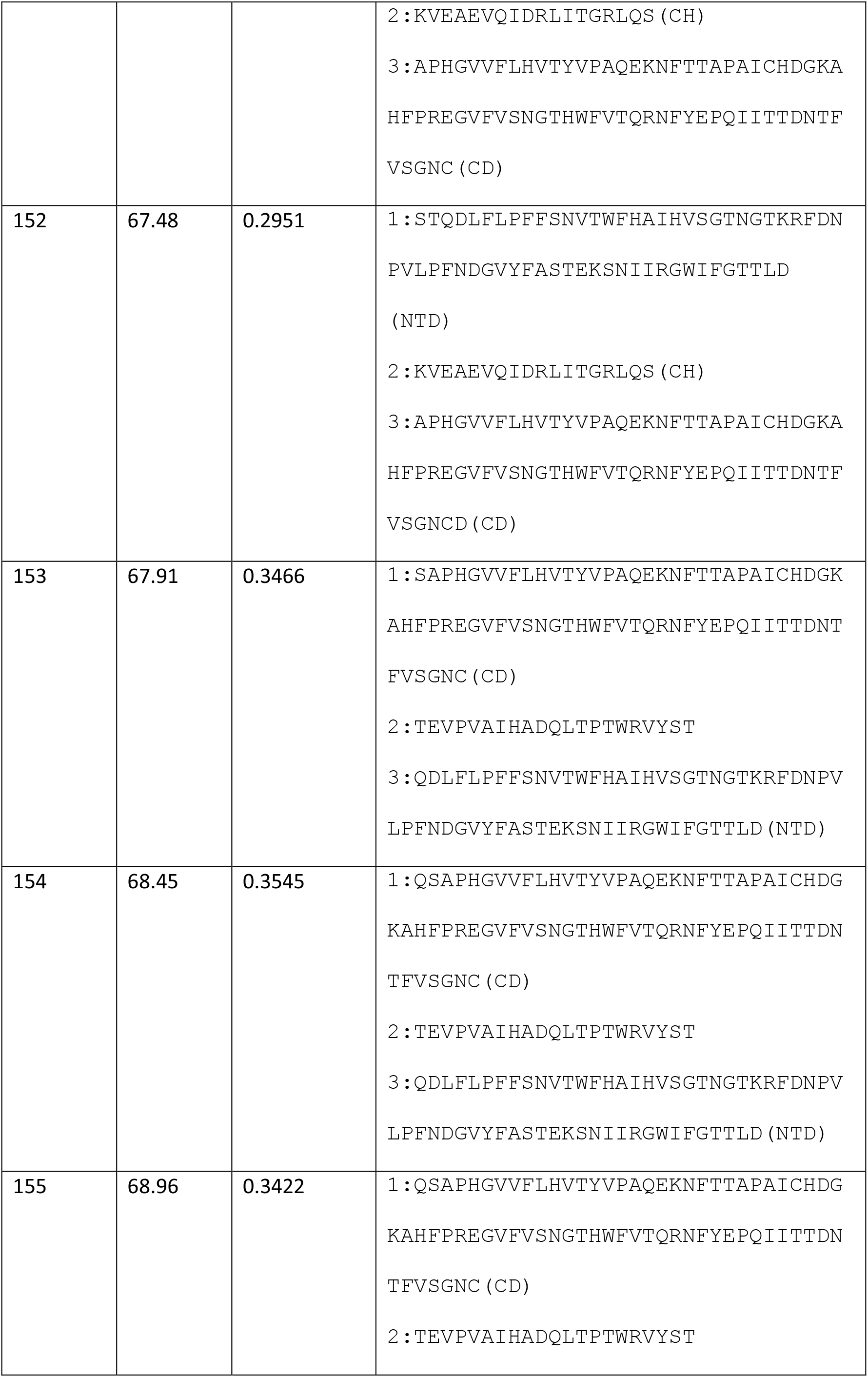
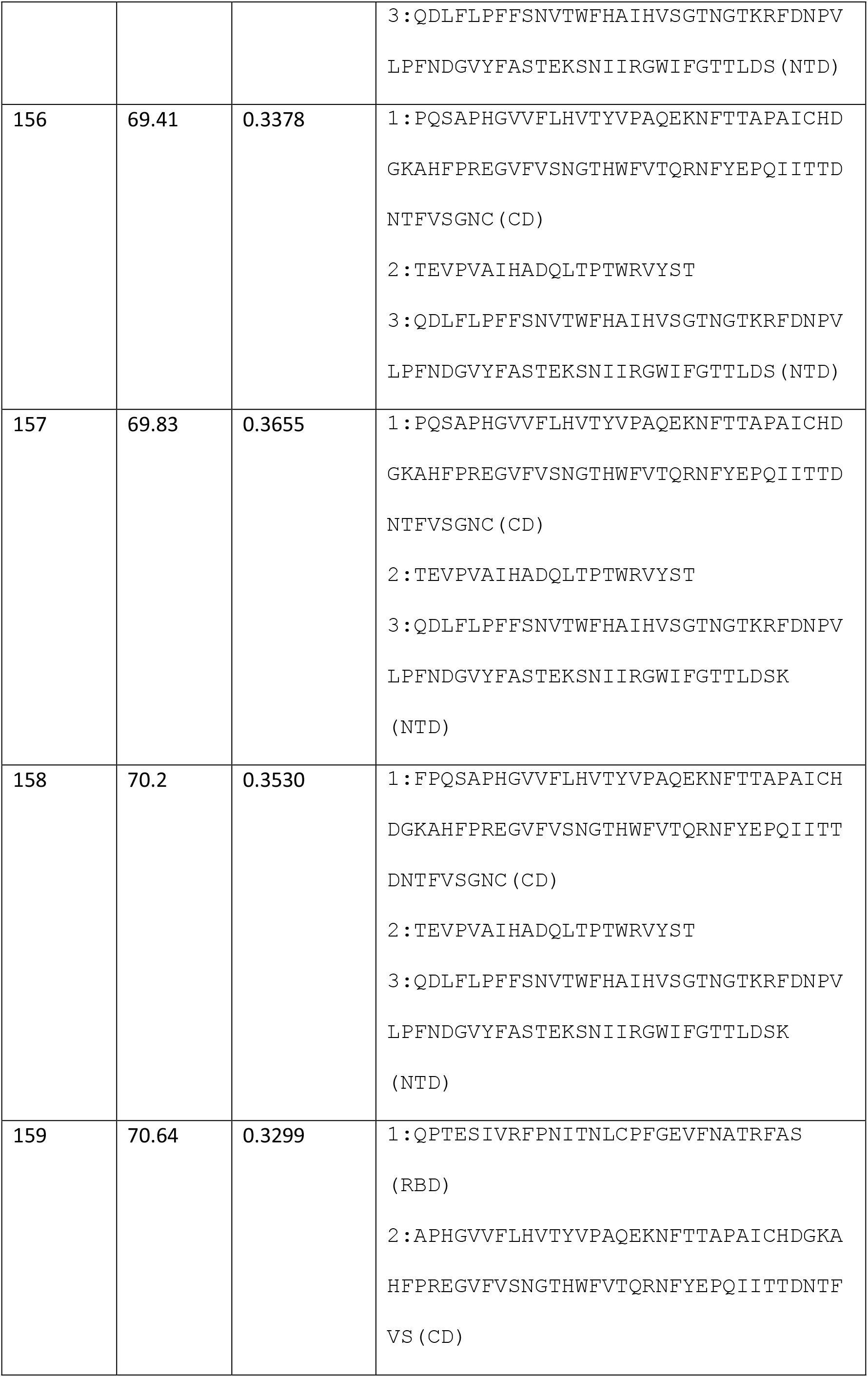
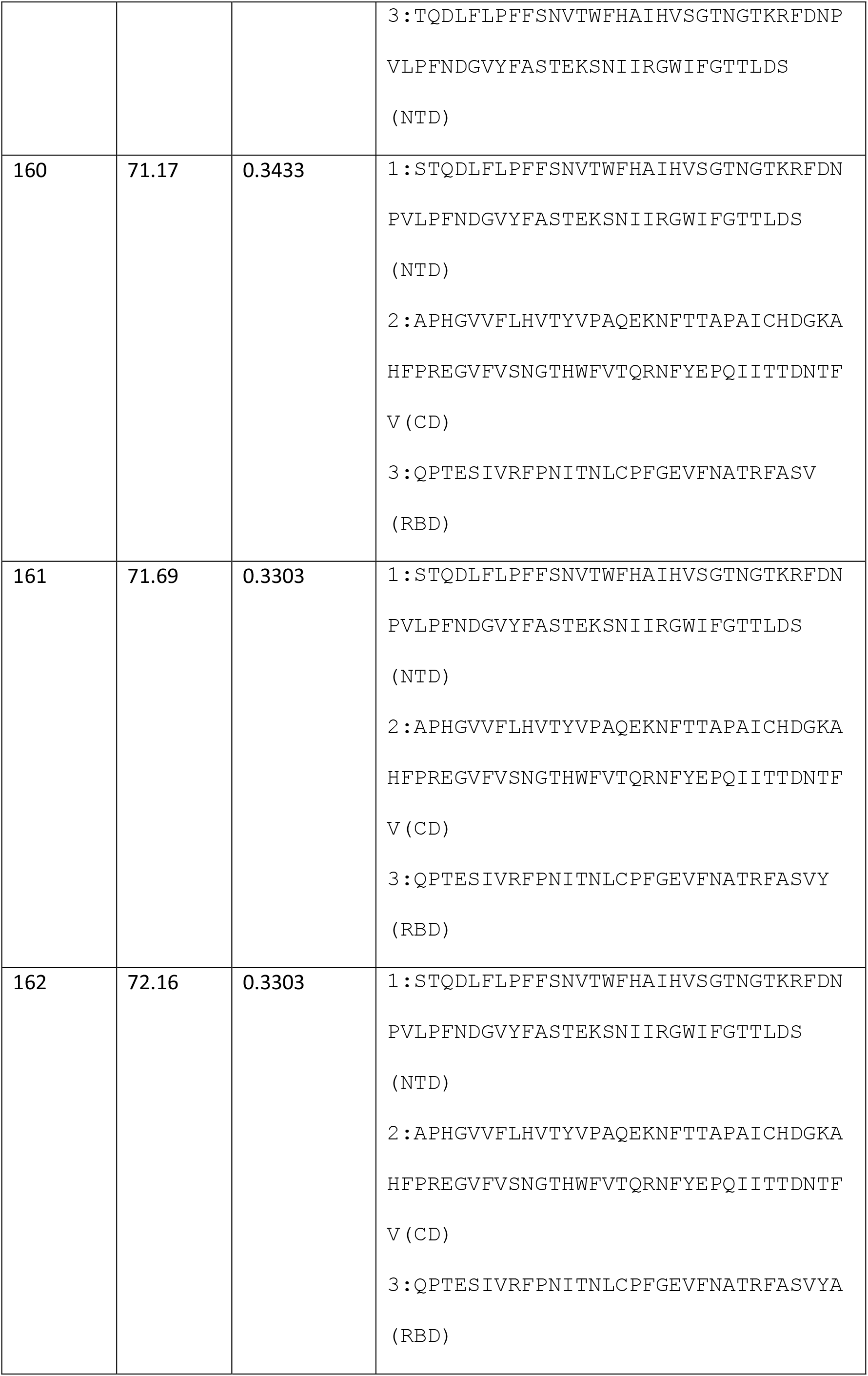
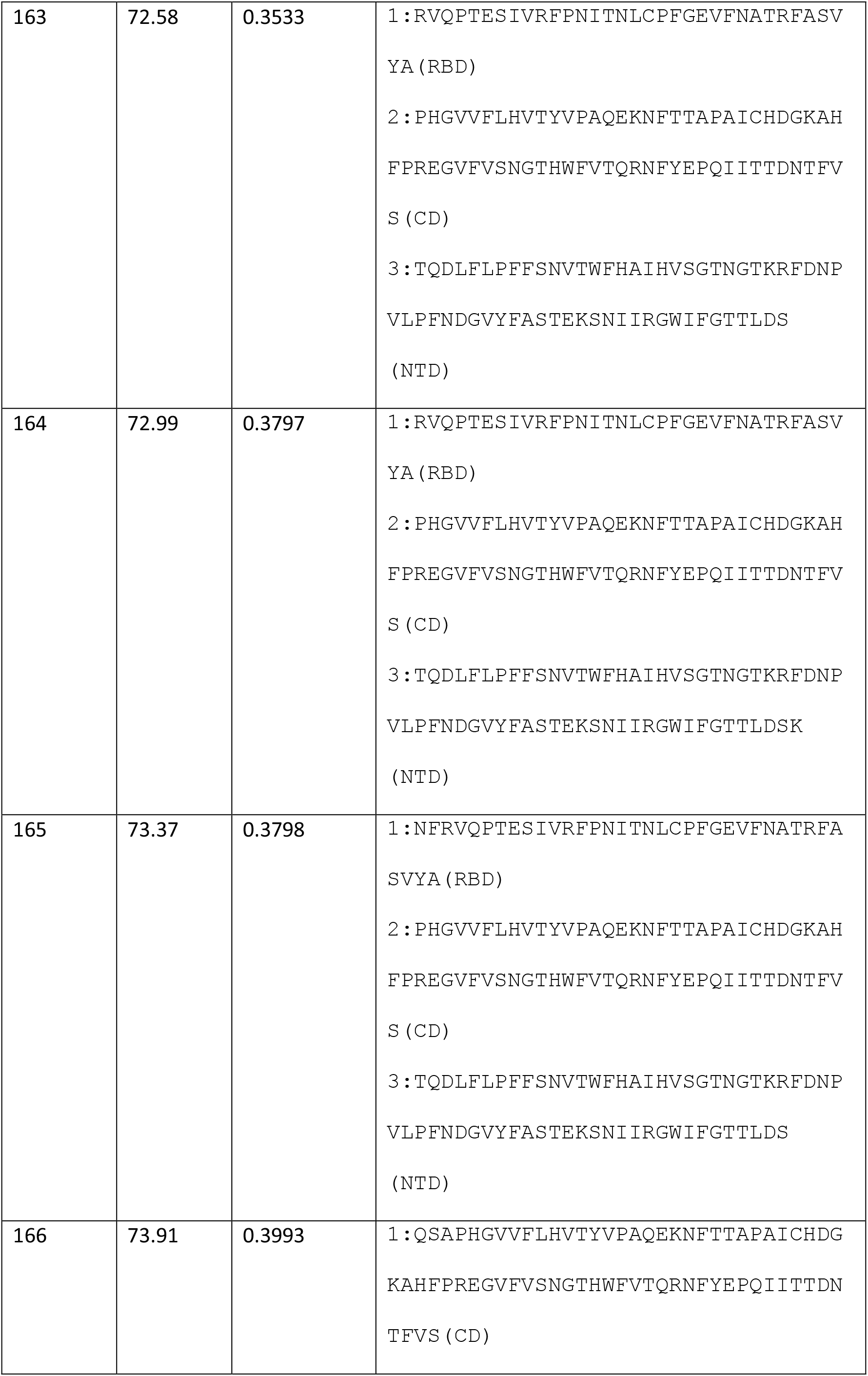
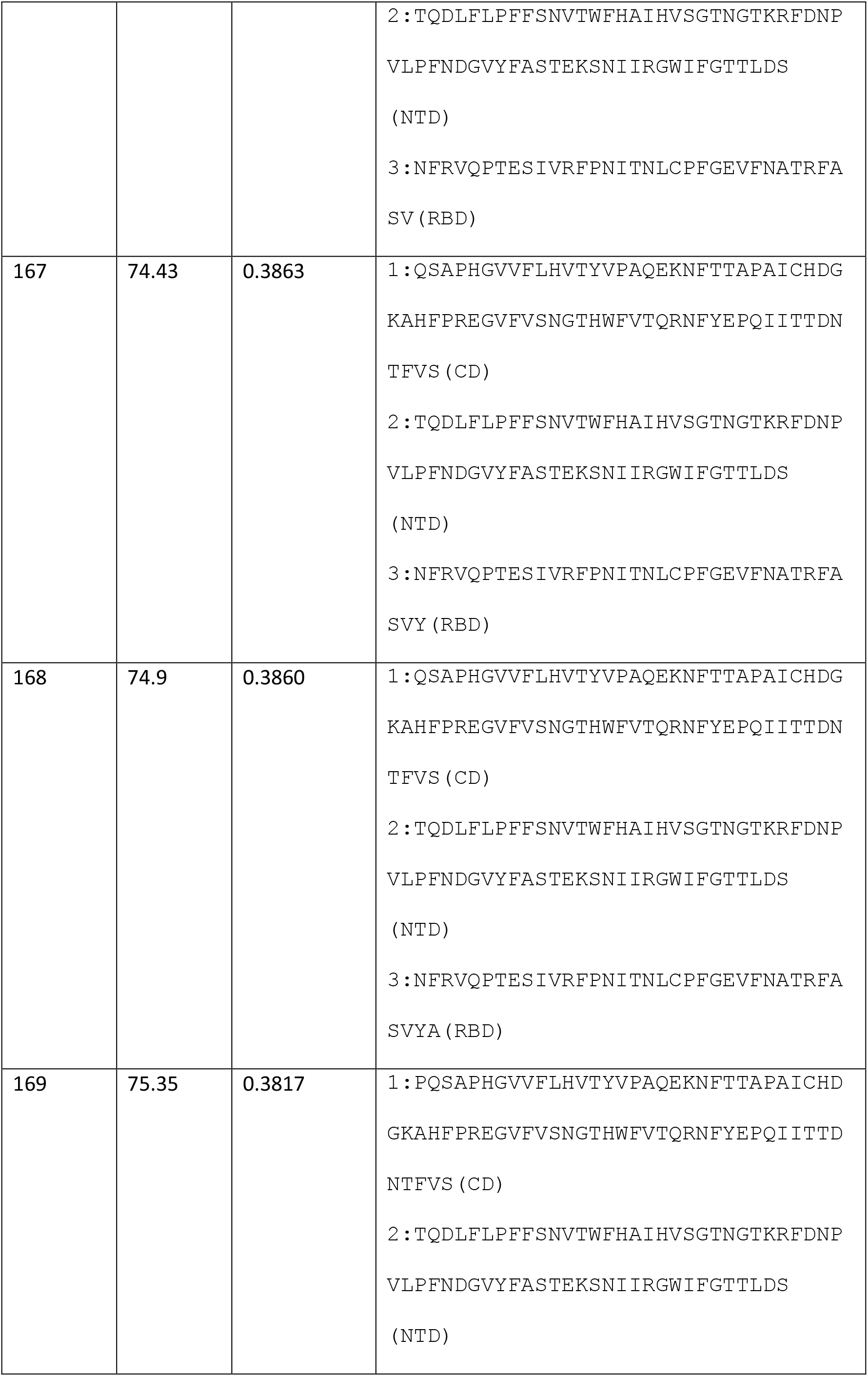
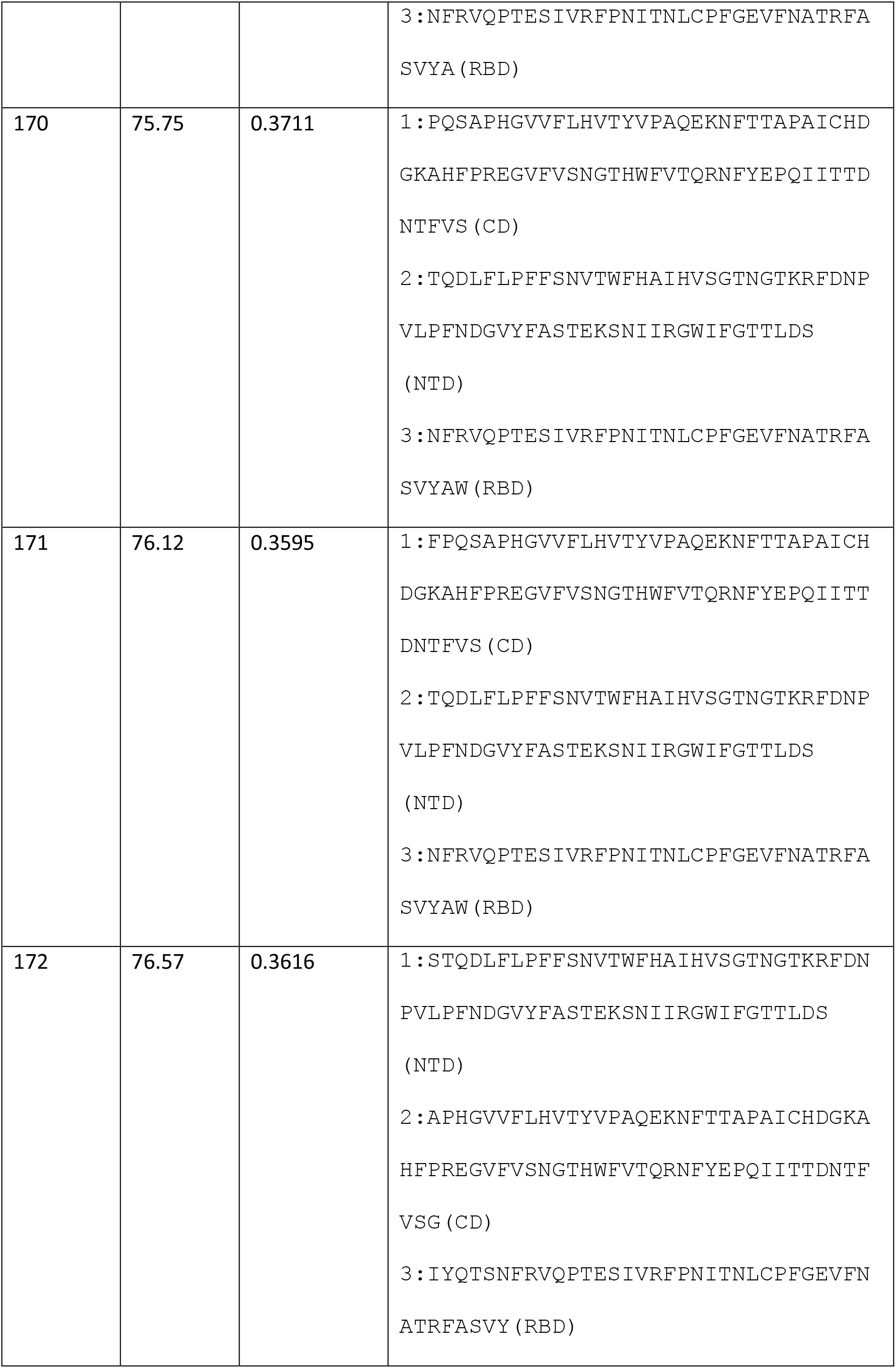
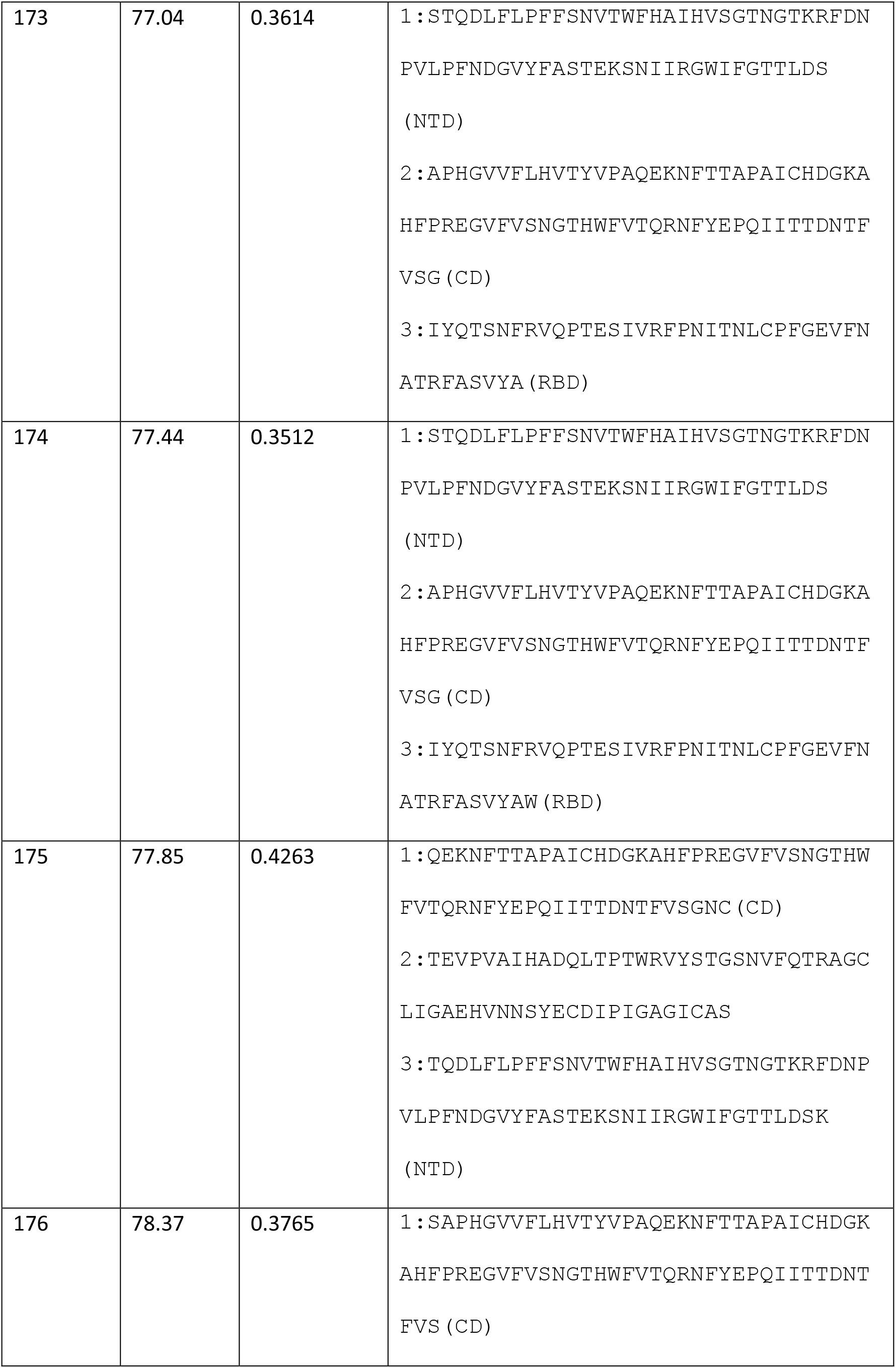
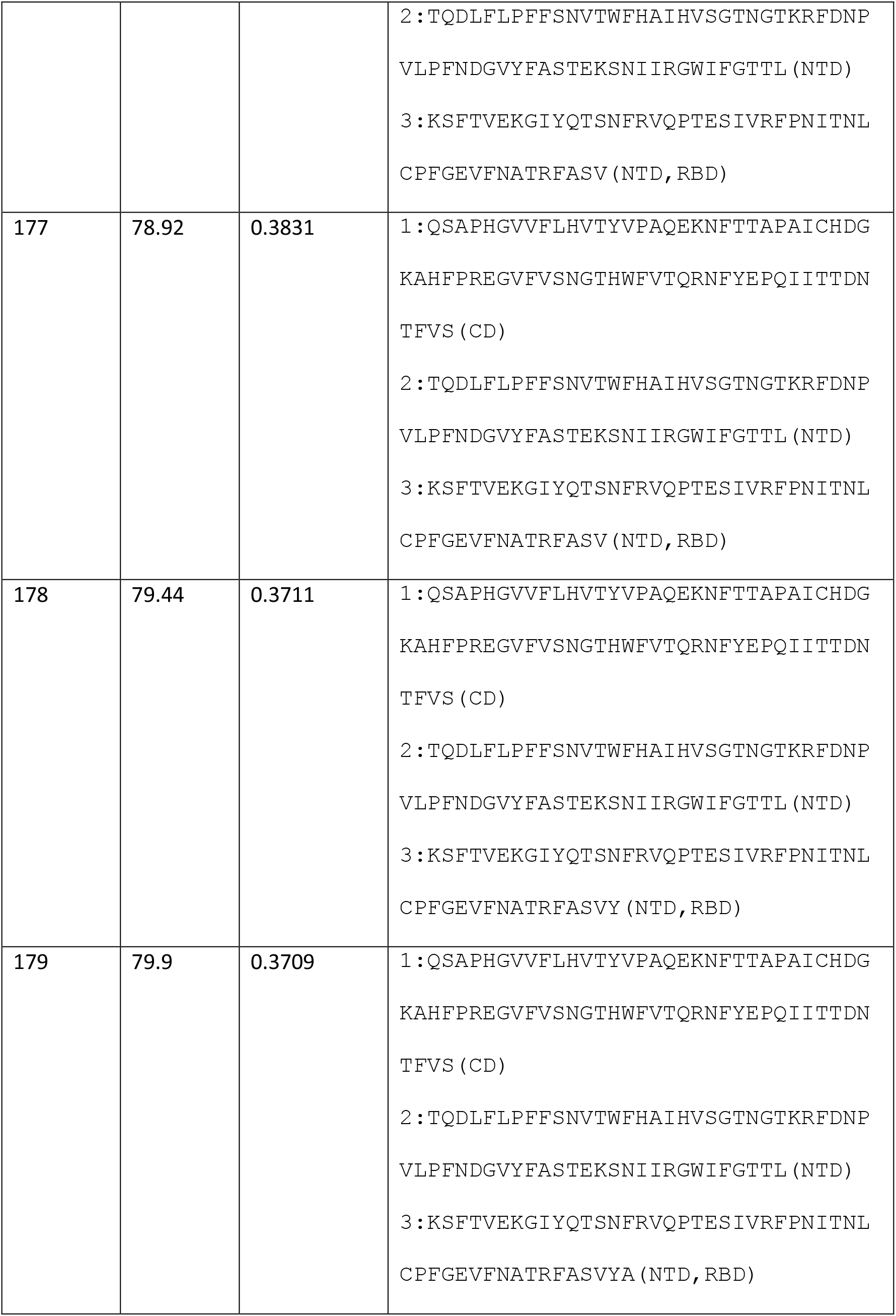
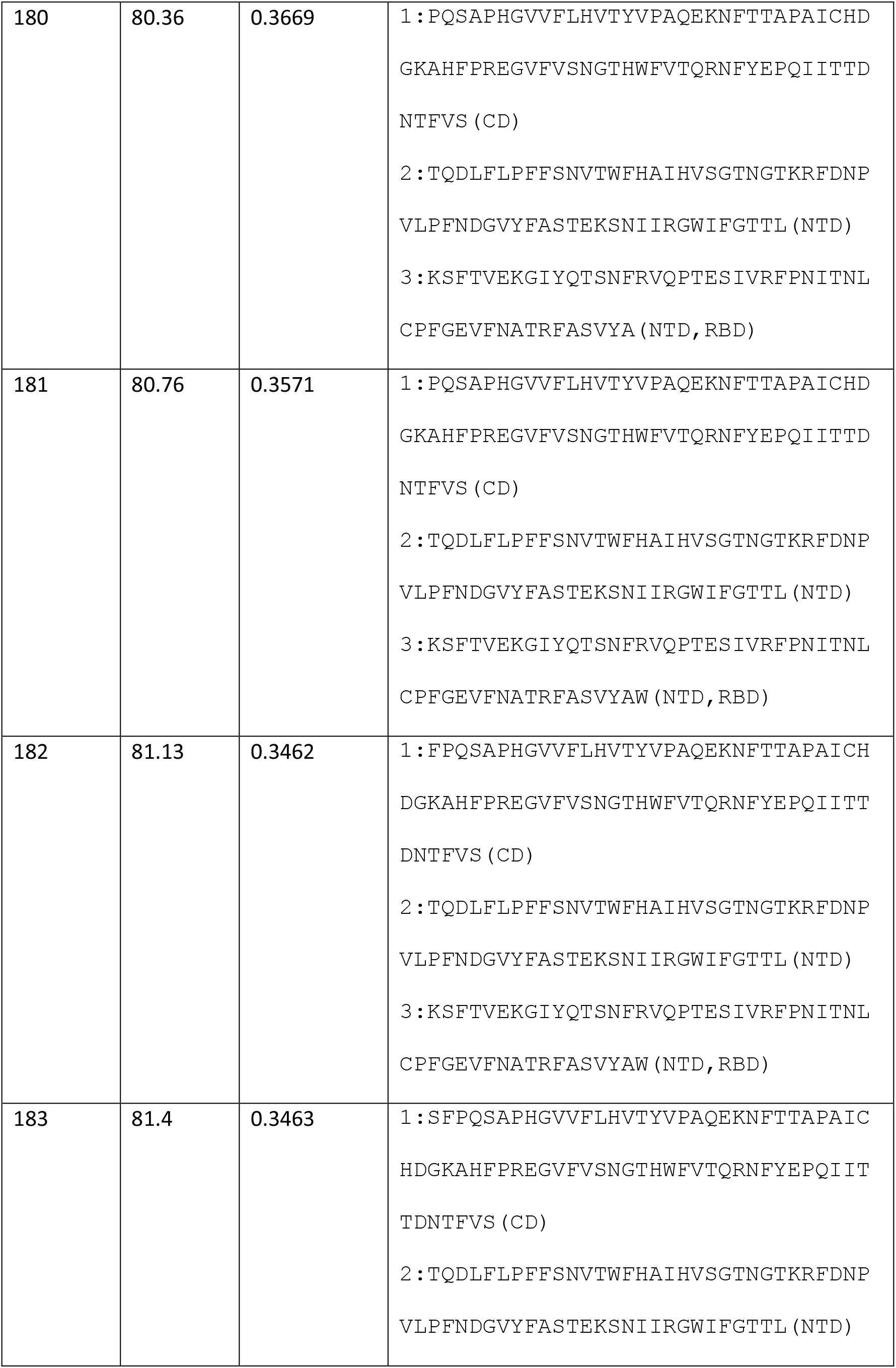
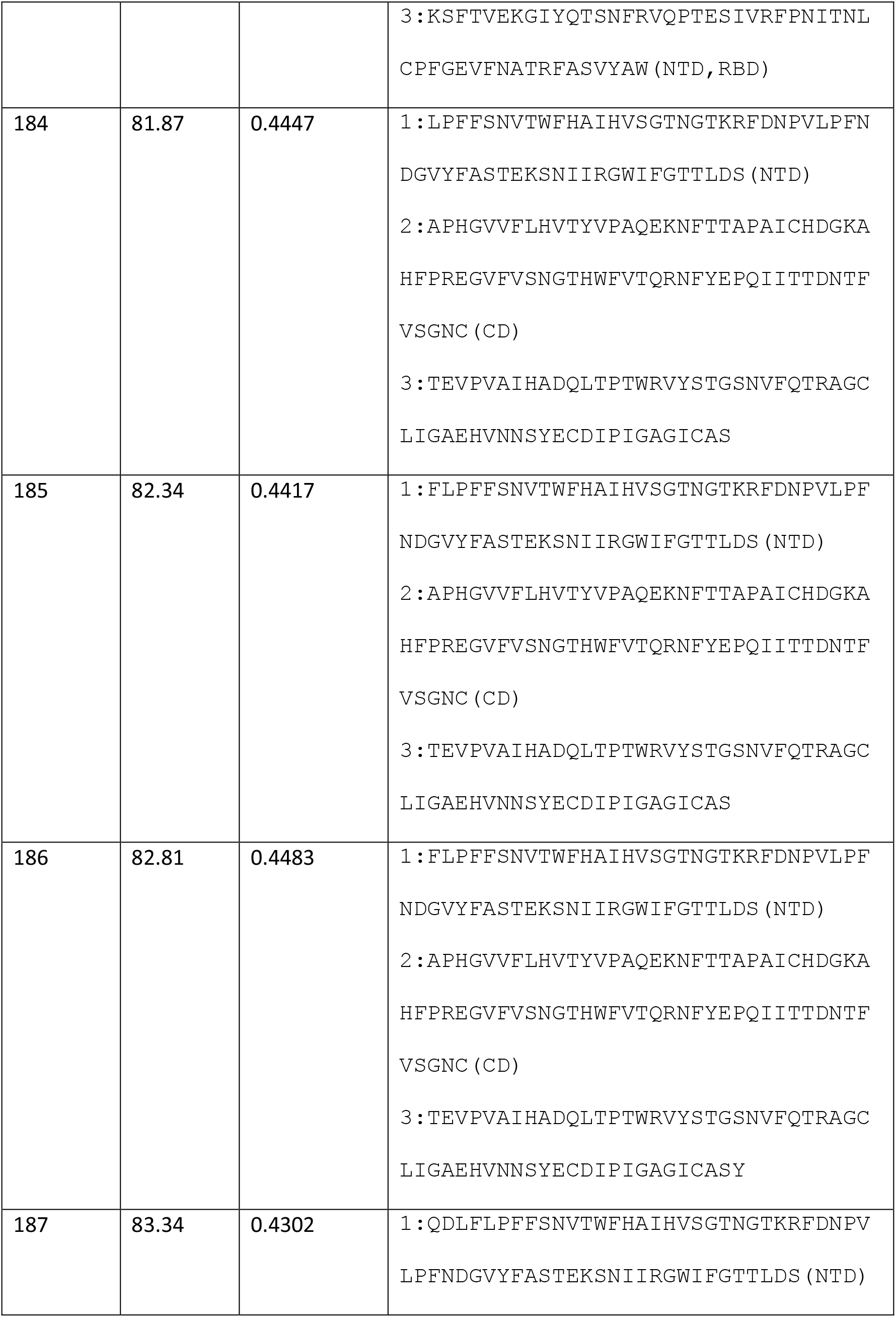
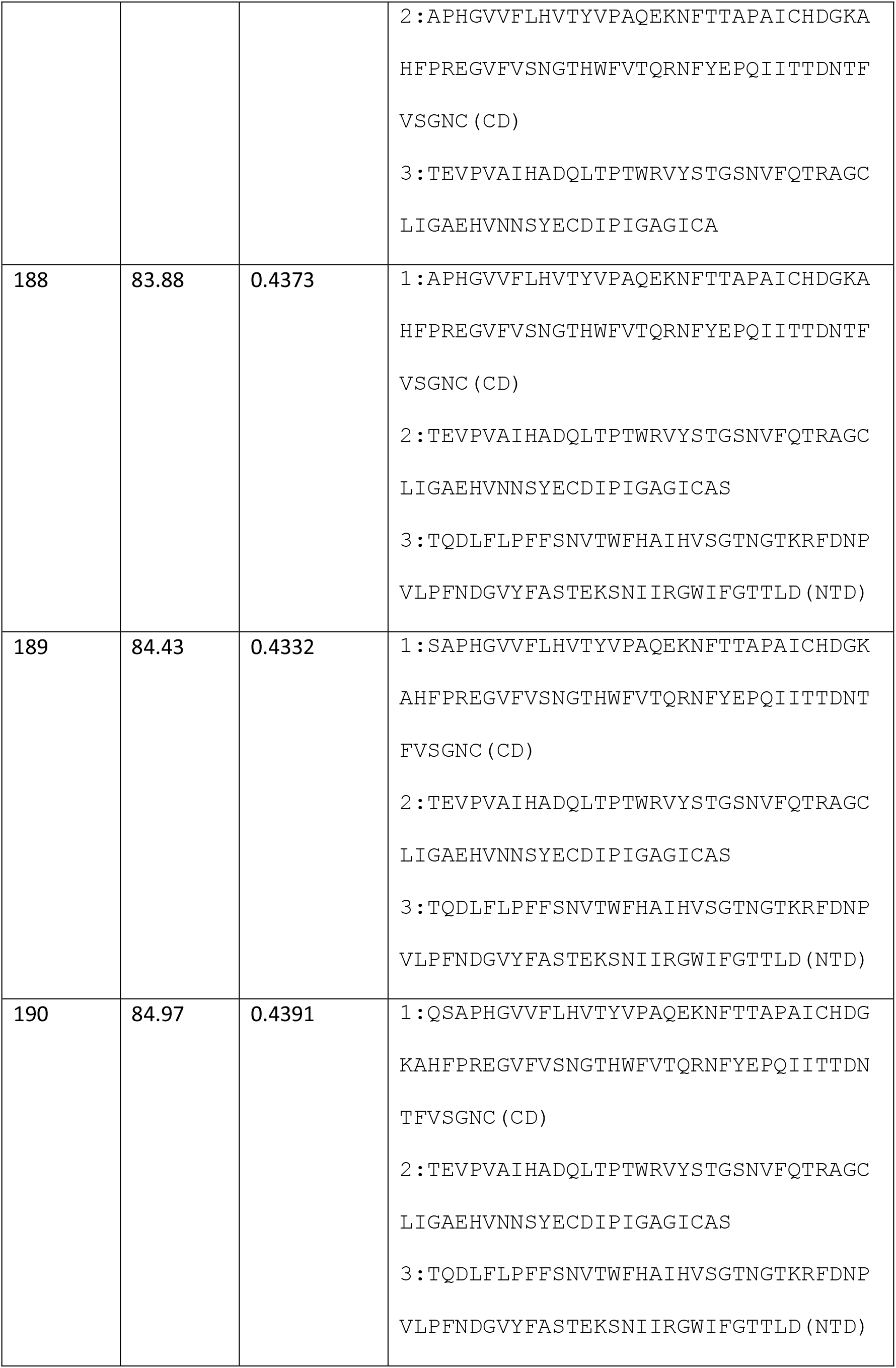
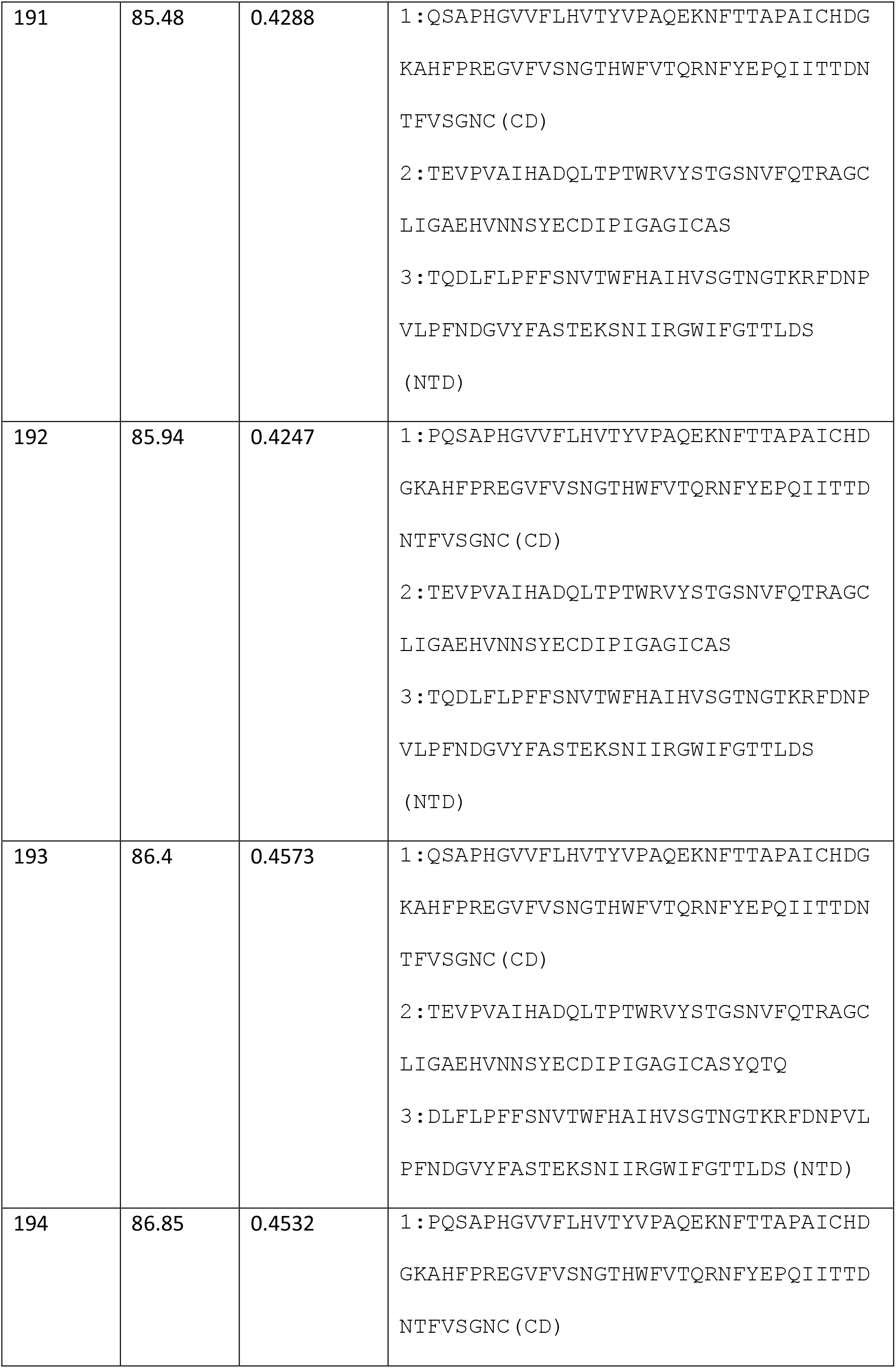
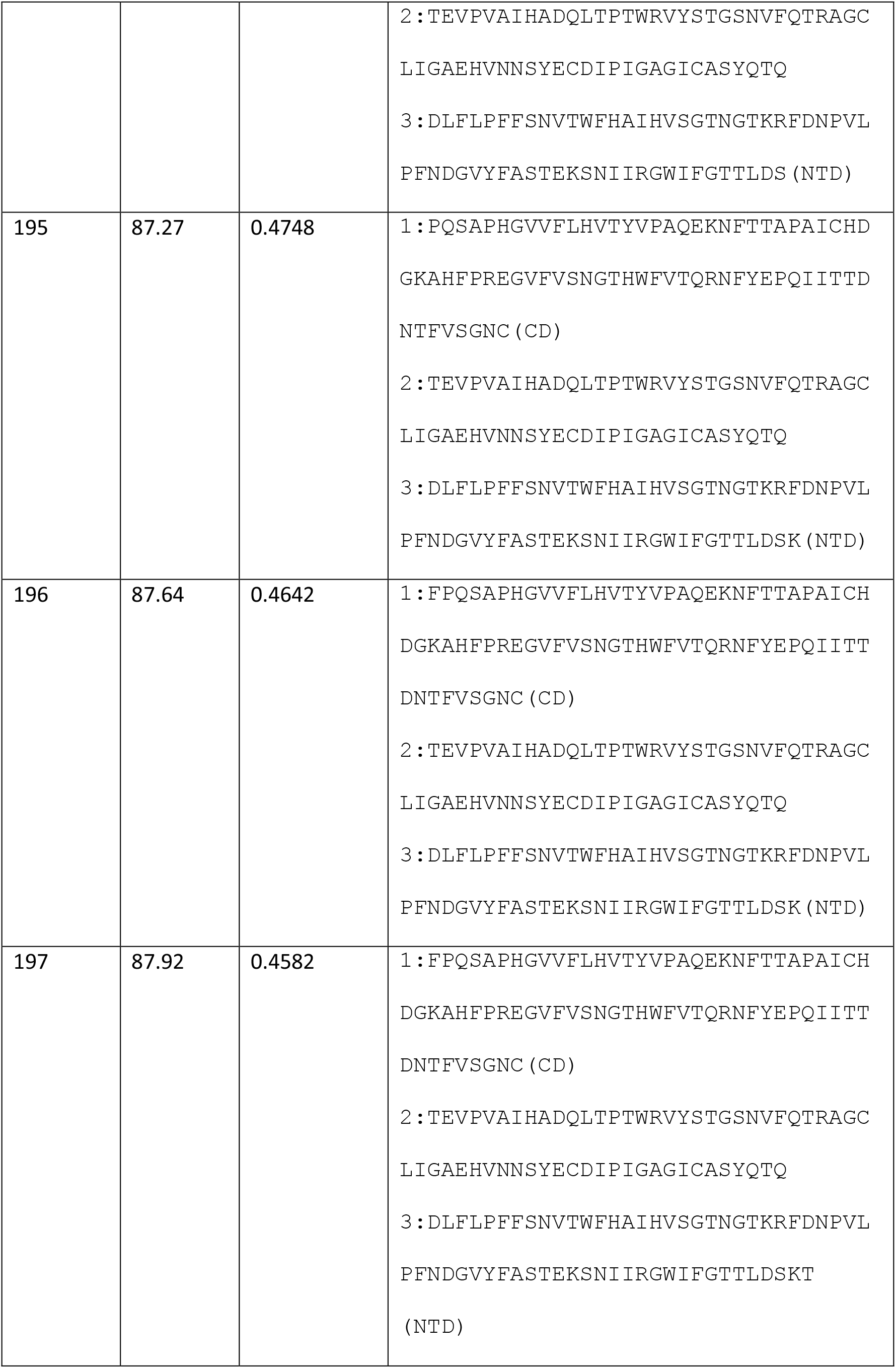
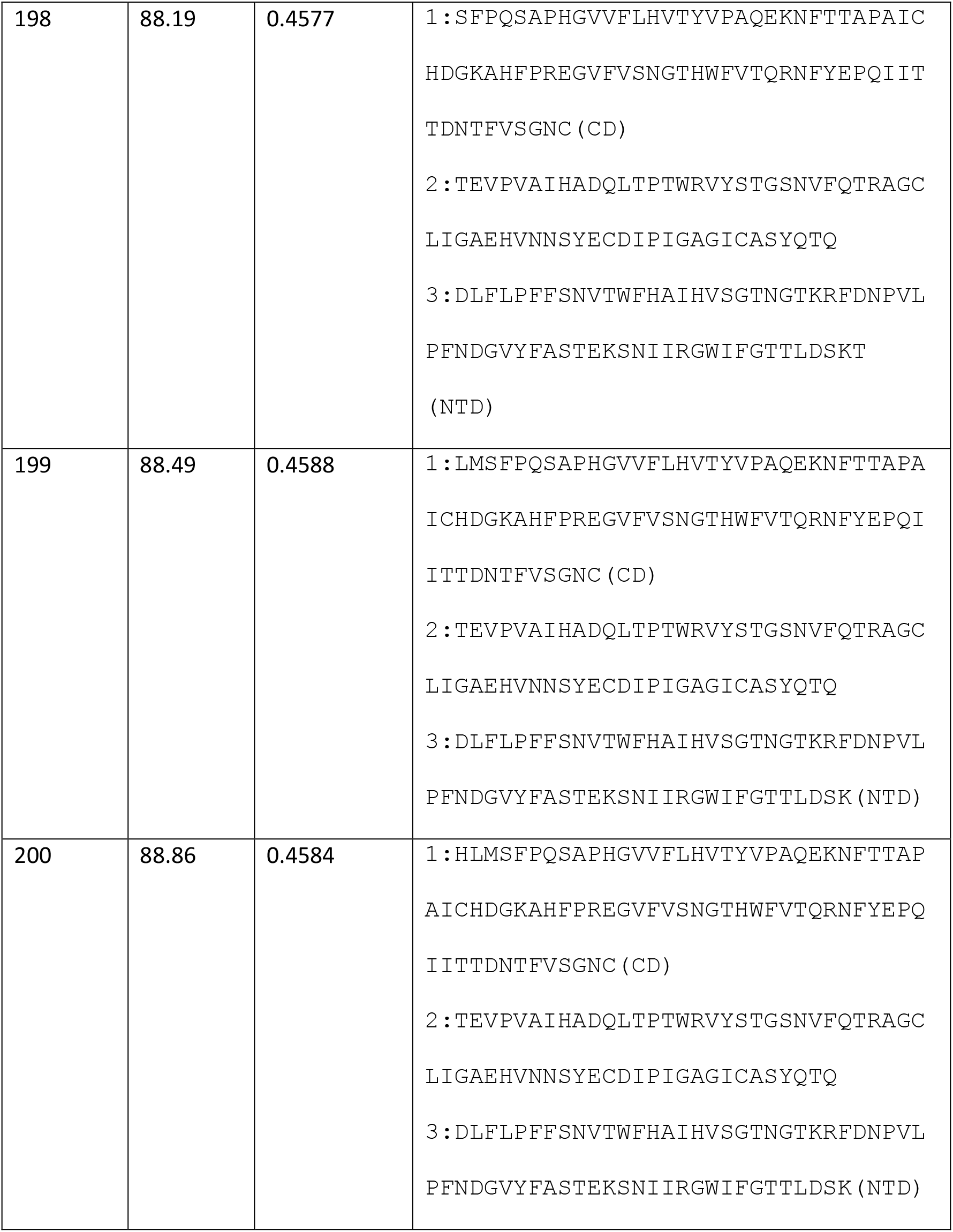
optimal weighted λ–superstring, λ values and VaxiJen overall prediction for lengths between 9 and 276.

We have chosen one of the candidate vaccines for which the overall prediction in VaxiJen exceeds 0.4, which is the threshold of the highest accuracy value beyond which the sequence is considered to be a probable antigen. Specifically, we have taken the peptide of length 22, STQDLFLPFFSNVTWFHAIHVS, which is contained in the NTD domain and has an overall prediction of 0.5545 in VaxiJen. We have synthesized the peptide, and we have done some in vivo experimental proofs to test its immunogenicity and putative efficiency.

First proof of concept regards to immunogenicity of the antigen prepared, SARS-CoV-2-NTD peptide of 22 amino acids (here named as COVID-19-NTD). We used a procedure previously described ([1]) that evaluates the best immunogenic epitopes for preparing vaccines. The assay consists of measuring the delayed type hypersensitivity (DTH) response of a vaccine vector, such as dendritic cells (DC) loaded with the peptide, in a vaccine formulation with an adjuvant. Next, we inoculated the vaccine formulation into the left hind footpads of mice, serving the non-inoculated right hind footpads as basal controls. Forty-eight hours later, we measured the DTH response as the swelling of the left hind footpads compared to the right hind footpads. We also included empty DC in these experiments and DC loaded with a control peptide non-related to SARS-CoV-2 virus. Analysis of DTH responses indicated that DC loaded with COVID-19-NTD peptide were significantly higher than immune responses elicited by DC loaded with the control peptide or empty DC (blue bars in Figure 1). Next, we collected the popliteal lymph nodes and cultured them *in vitro* with 1 μg/mL of each peptide, COVID-19-NTD, control peptide or saline for 72 hours to examine the main immune populations by flow cytometry. We observed that the highest percentages of immune cells corresponded to CD19^+^ cells (27,63 %) that usually correspond to B cells, followed by MHC-II^+^ cells (20, 45%) that usually labels DC and macrophages, next CD4^+^ T cells (10,39%) and CD8^+^ T cells (4,61%). The control peptide does not produce significant immune responses, we only observed very small percentages of CD19^+^ cells (4,61%) and moderate ranges of MHC-II^+^ cells (11,5%) (DC-peptide CONT bars in Figure 1). Empty DC did not induce significant percentages of immune cells. These results indicate a clear induction of immune cells by DC vaccines loaded with COVID19-NTD peptide that stimulate immune cells involved in antibody formation, such as B cells, DC and CD4^+^ T cells. While not predominant, cytotoxic immune responses caused by CD8^+^ T cells are also induced by DC vaccines loaded with COVID-19-NTD peptide. These results are not surprising since CD4+ and CD8+ T cell epitopes are recovered from patients with mild and severe COVID-19 that were specific for the spike protein ([8]).

**Figure 1.**
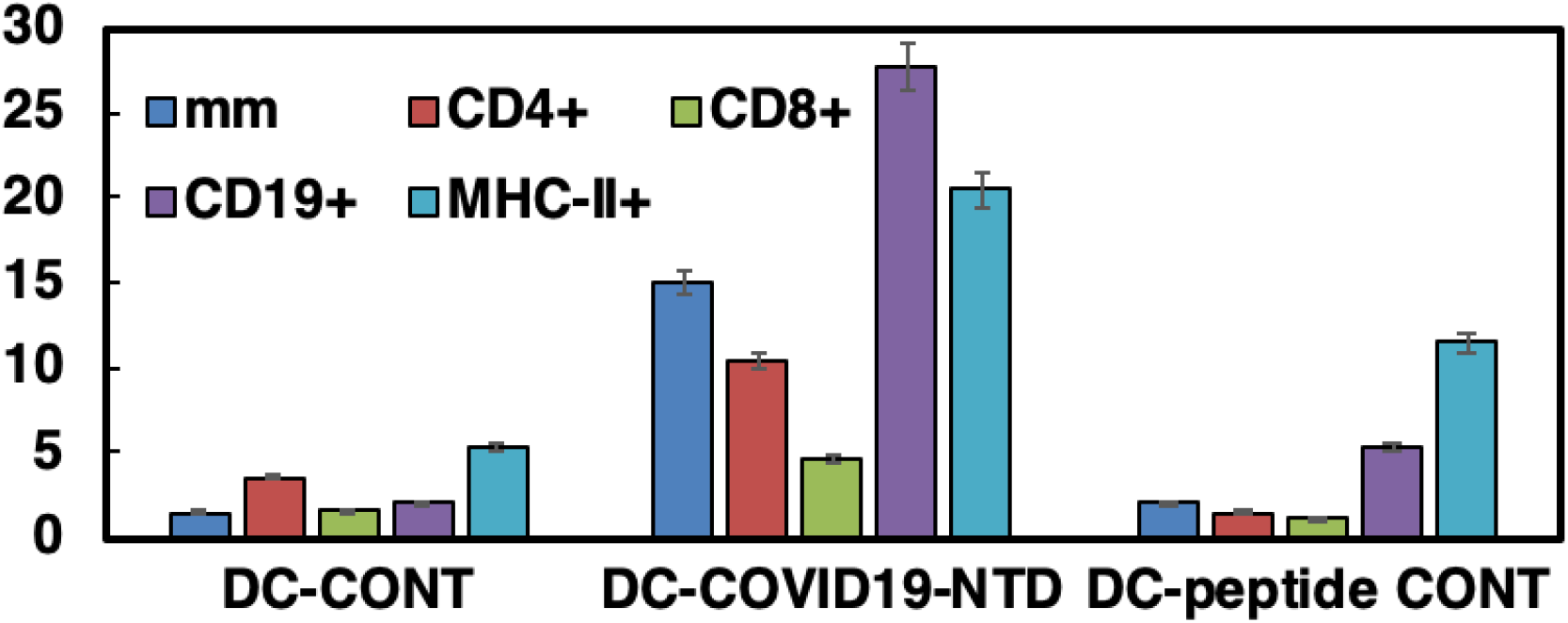
Immunogenic abilities of COVI19-NTD peptide in vaccine platforms. Mice (C57BL/6, n = 5) hind footpads are inoculated with the DC vaccines (10^6^ cells/mice) loaded with the different peptides (COVID-19-NTD or peptide control) or remained as empty DC in formulations with the adjuvant DIO-1 (40 ng/mL). 48 hours later, footpad swelling was measured with a caliper (dark blue bars) and expressed as the differences in mm between left and right hind footpads. Results are the mean ± SD of three different experiments (*P* < 0.05). Popliteal lymph nodes are next isolated from mice legs and after homogenization, immune cell populations are analyzed by flow cytometry. Percentages of CD4^+^ (red bars), CD8^+^ T cells (green bars), CD19+ (B cells, purple bars) and MHC-II+ positive cells, mainly DC or macrophages (light blue bars) are shown. Results are expressed as the percentages of positive cells ± SD of three different experiments (P < 0.05).

Second proof of concept related with the production of cytokines, either pro-inflammatory cytokines such as TNF-a, IFN-g, IL-2, KC and IL-12 or anti-inflammatory cytokines as IL-4, IL-6, MIP-2 or IL-10. Covid-19 cytokine storm in severe patients correlates with very high levels of TNF-a, IL-6, IL-4 and IL-10, as well as with a clear deficiency in the production of IFN related cytokines (*i.e,* IFN-a, IFN-g or IL-12) ([9]). Our results in Figure 2 showed that DC loaded with COVID19-NTD peptide produced mainly the pro-inflammatory cytokines, IL-12, IL-17A and IL-2. However, this DC-COVID-19-NTD vaccine platform does not induce anti-inflammatory cytokines participating in COVID-19 cytokine storm such as IL-6, IL-10 or TNF-a (bars labelled with DC-COVID19-NTD in Figure 2). Interesting, DC-COVID-19-NTD vaccines induced high levels of IFN-g (blue bars in Figure 2) and barely no levels of MIP-2, an inflammatory cytokine that recruits inflammatory macrophages (grey bars in Figure 2). The lack of significant levels of IL-4 (orange bars in Figure 2), while high levels of IL-2 (red bars) suggests induction of a Th1-Th17 type immune responses, but with no exacerbation of the pro-inflammatory cytokines as TFN-a or MIP-2. In summary, high levels of IFN-g and especially IL-12 cytokines that involved on vaccine efficiency and anti-viral responses, prompted us to suggest that COVID19-NTD peptide might function as an immunogenic epitope. COVID19-NTD peptide might be a good candidate to prepare efficient vaccine platforms that induce not only antibody production but strong anti-viral T cell responses.

**Figure 2.**
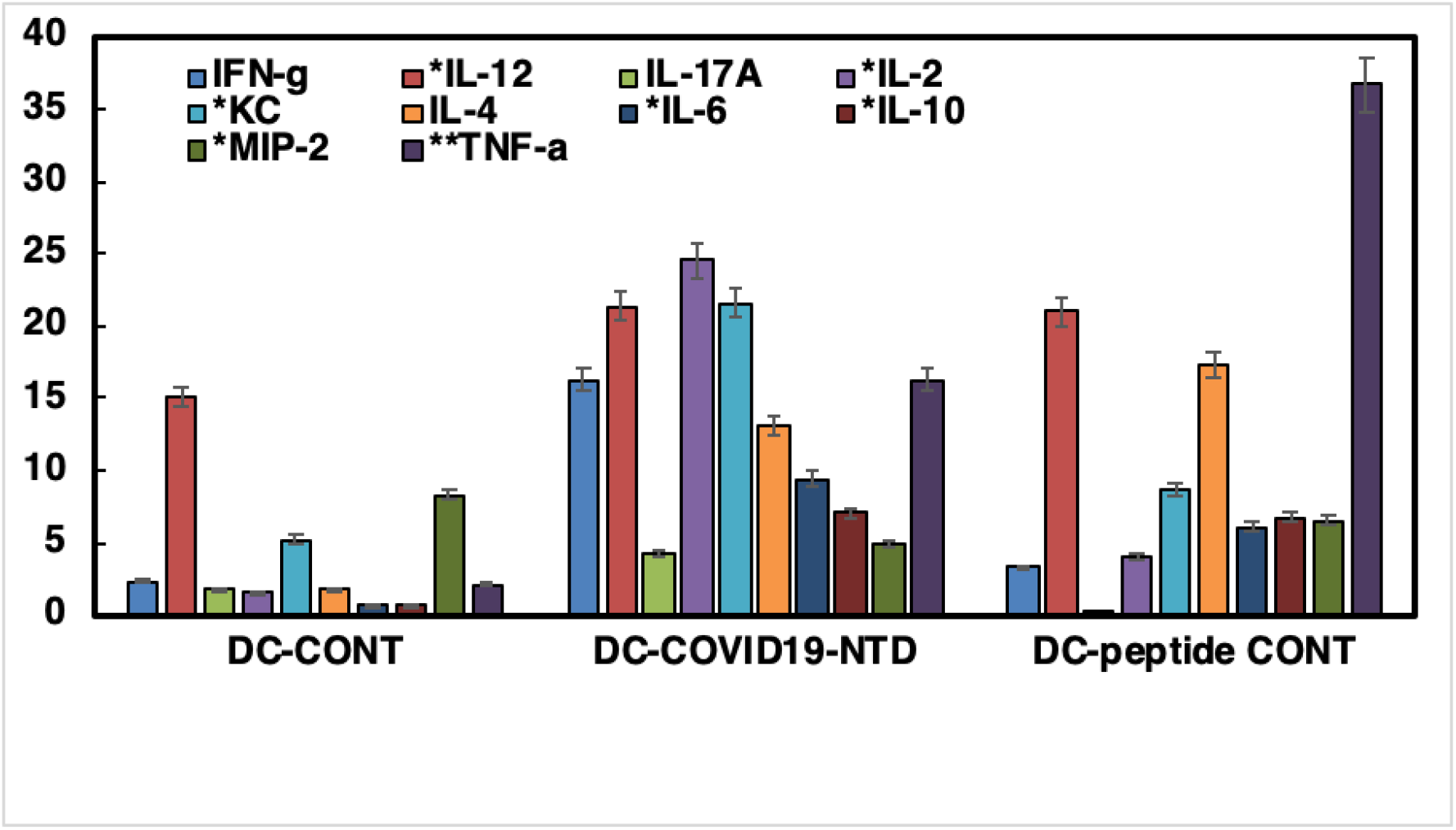
Cytokine levels of mice inoculated with DC vaccine platforms. Cytokine levels detected in sera of mice as in Figure 1 and measured with a multiparametric Luminex kit from Merck. Results are expressed as pg/mL of each cytokine ± SD of triplicate samples (*P* ≤ 0.5).

## 4. Discussion

The selected candidate vaccine elicited a strong immune response in mice with Th1-Th17 pro-inflammatory features with strong stimulation of cells involved in antibody production but also with production of anti-viral cytokines. This shows the adequacy of using λ-superstrings to find an efficient vaccine against SARS-CoV-2.

The exploration of the candidate vaccines in the table of the previous section could lead to the obtention of a vaccine that covers a high percentage of the population. Candidate vaccines in the table for which the overall prediction in VaxiJen is greater than 0.4, which is the threshold for the highest accuracy, are specially interesting, but it could be useful to consider also values close to 0.4, say for instance over 0.37.

